# Clostridia isolated from helminth-colonized humans promote hatching of *Trichuris muris*

**DOI:** 10.1101/2022.02.10.479991

**Authors:** Shushan Sargsian, Ze Chen, Soo Ching Lee, Amicha Robertson, Julia Sproch, Joseph C. Devlin, Mian Zi Tee, Yi Xian Er, Richard Copin, Adriana Heguy, Alejandro Pironti, Victor J. Torres, Kelly V. Ruggles, Yvonne A.L. Lim, P’ng Loke, Ken Cadwell

## Abstract

Soil transmitted intestinal worms known as helminths colonize over 1.5 billion people worldwide. Although helminth colonization has been associated with altered composition of the gut microbiota, such as increases in Clostridia, individual species have not been isolated and characterized. Here, we isolated and sequenced the genome of 13 Clostridia from the Orang Asli, an indigenous population in Malaysia with high prevalence of helminth infections. Metagenomic analysis of 650 fecal samples from urban and rural Malaysians revealed higher prevalence and abundance of these isolates compared to individuals in the United States, with *Peptostreptococcaceae* family members displaying a specific association with helminth colonization. Remarkably, *Peptostreptococcaceae* isolated from the Orang Asli displayed superior capacity to induce hatching of eggs from the murine helminth *Trichuris muris*. These findings support a model in which helminths select for gut colonization of microbes that in turn support their life cycle by promoting egg hatching.

## Introduction

Intestinal parasitic worms known as helminths affect roughly 24% of the world population (WHO, 2020). The most common species that infect humans are the whipworm *Trichuris trichiura*, roundworm *Ascaris lumbricoides*, and hookworms *Necator americanus* and *Ancylostoma duodenale*, which are associated with maladies ranging from diarrhea to stunted growth and malnutrition (Bethony et al., 2006; WHO, 2020). Although helminths cohabitate the mammalian gastrointestinal tract alongside trillions of microbes and can influence the diversity and composition of the microbiome (Kupritz et al., 2021), species level resolution and functional characterization of helminth-associated bacteria has lagged behind. Microbiome studies have been dominated by those examining individuals in industrialized nations where the prevalence of helminths is low. Neglecting to study helminth-associated microbes may result in a lack of not present in the microbiomes of better characterized populations (Blaser, 2016; Blaser and Falkow, 2009). Furthermore, medical advances gained from microbiome studies may be biased towards individuals in high income countries and insufficient in addressing the healthcare needs of marginalized groups in low- and middle-income countries.

The microbiome of helminth colonized individuals display cohort-specific compositions (Cooper et al., 2013; Easton et al., 2019; Jenkins et al., 2017; Kay et al., 2015; Lee et al., 2014; Martin et al., 2018; Rosa et al., 2018). Inconsistent associations between microbes and helminth colonization likely reflects the unique ethnic populations, environment, and parasites associated with different geographic regions, highlighting the importance of deeper characterization of population-specific microbiomes. We previously reported higher gut microbial diversity in the Orang Asli, an indigenous population in Malaysia with a high prevalence of helminth infections, especially *T. trichiura*, compared to nearby urbanized Kuala Lumpur (Ramanan et al., 2016). Following deworming treatment with the anthelmintic drug albendazole, the microbiomes of the Orang Asli showed decreased alpha-diversity accompanied by increased Bacteroidales and decreased Clostridiales. Colonization of mice with *Trichuris muris*, the *Trichuris* species that infects mice, or the roundworm *Heligmosomoides polygyrus bakeri* reveal similar shifts in the microbiome that support a positive relationship between helminth colonization and species belonging to the Clostridiales order (Ramanan *et al.*, 2016; Zaiss et al., 2015).

Clostridia are a polyphyletic class of Firmicutes that includes pathogenic and commensal Clostridiales species with ill-defined taxonomic relationships sharing the key feature of being spore-forming obligate anaerobes (Dürre, 2014; Nagano et al., 2012). Clostridia have received considerable attention as immunomodulatory members of the microbiota due to their metabolic activity and capacity to confer colonization resistance against pathogens (Atarashi et al., 2013; Becattini et al., 2017; Caballero et al., 2017; Kim et al., 2017; Sokol et al., 2008; Stefka et al., 2014). Expansion of Clostridia during helminth colonization mediates amelioration of disease in mouse models of inflammatory bowel disease and asthma (Ramanan *et al.*, 2016; Zaiss *et al.*, 2015), supporting the importance of genomic and functional characterization of helminth-associated bacteria in humans.

Intestinal bacteria promote gastrointestinal colonization by helminths, indicating bidirectional communication between parasites and the microbiome (Lawson et al., 2021; Reynolds et al., 2014). For example, bacteria promote hatching of *T. muris eggs*, and *T. muris* cannot complete its life cycle in germ-free mice that lack a microbiome (Hayes et al., 2010; Venzon et al., 2021; White et al., 2018). It is unclear whether specific bacterial species display differential effects on helminth reproduction. Here, we report the genomic characterization of bacteria isolated from helminth-colonized Orang Asli villagers. Our findings identify specific taxa, especially those belonging to the *Peptostreptococcaceae* family, that are uniquely associated with helminth colonization and promote *T. muris* egg hatching, providing evidence of parasite-microbiome co-adaptation.

## Results

### Isolation and identification of spore-forming Firmicutes from helminth-colonized individuals

Our previous analyses of stool specimens from the Orang Asli using 16S sequencing showed a positive correlation between *T. trichiura* burden and relative abundance of Clostridiales (Ramanan *et al.*, 2016). Therefore, we screened stool from 19 Orang Asli (9 female, 10 male, ages 5-51, *Trichuris* prevalence = 52.63%) from the Pangsun village to identify individuals with high *T. trichiura* burden to increase the probability of isolating Clostridiales species of interest (Table S1). We applied a previously described procedure in which fecal samples are treated with chloroform and passaged in germ-free mice to enrich for spore-forming bacteria (Atarashi *et al.*, 2013), to specimens from the Orang Asli individuals with the four highest *Trichuris* burdens. Isolates from samples #88 and #135 were discarded due to contamination during the procedure. We proceeded with isolating bacteria from the remaining two samples (#20110 and #762) with Kato Katz intensities of 4,375 and 420 eggs (Table S1, Figure S1A). From 75 initial colonies, we used 16S sequencing to identify 14 isolates with unique sequences and/or morphologies, designated OA1-14. Bacterial colonies displayed distinct visual properties including mucoid and filamentous morphologies and hemolytic activity, providing evidence that they represent non-redundant isolates (Figure 1A).

**Figure 1.**
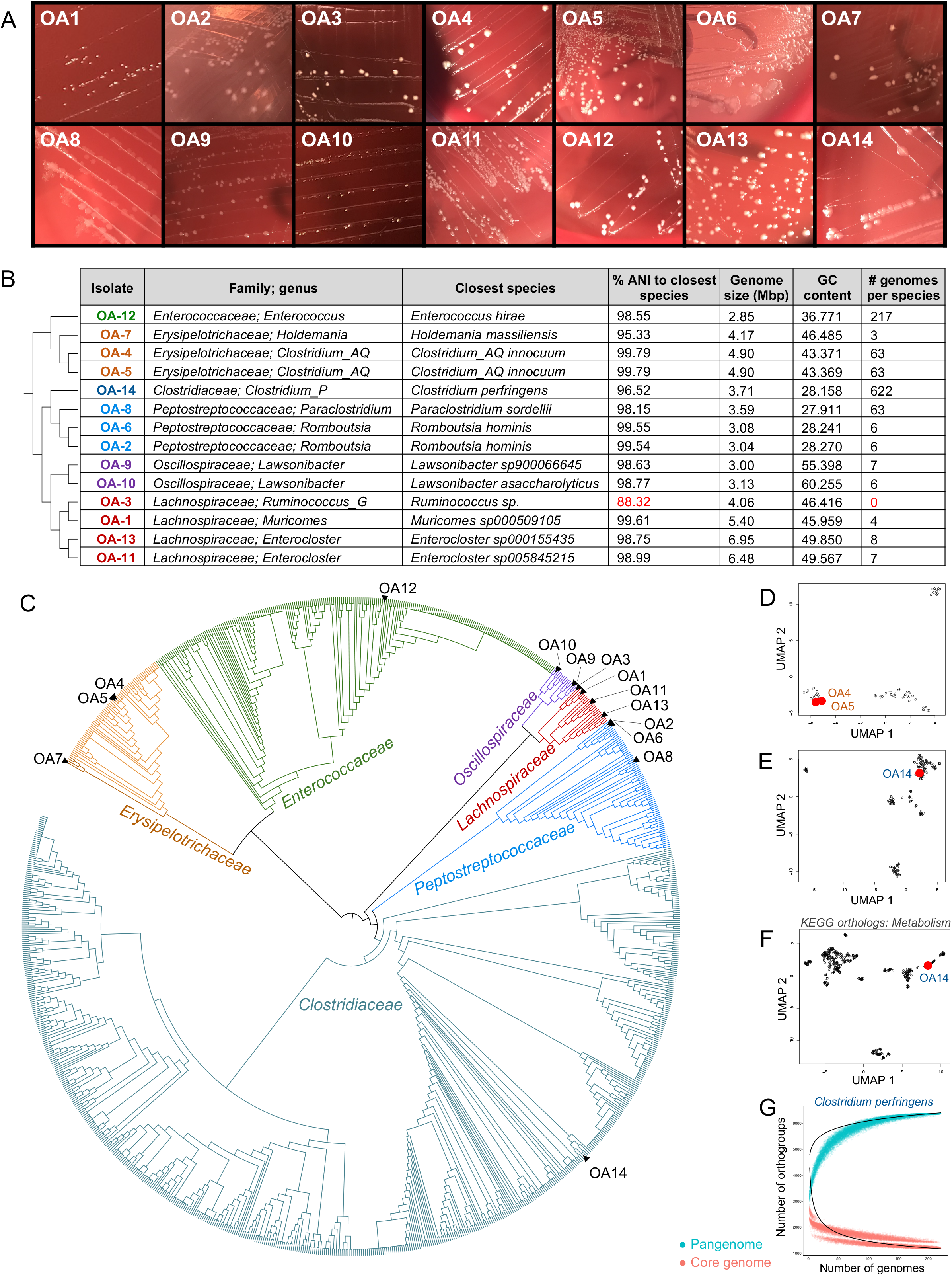
Isolation and identification of spore-forming Firmicutes from helminth-colonized individuals. (A) Representative images of OA isolates 48 hours after streaking on brucella blood agar. (B) Taxonomic identities and genomic features of OA isolates based on full genome sequences. Closest known relative refers to species with the highest average nucleotide identity (ANI). Red: denotes new species with no matches 95% ANI. The phylogenetic tree to the left of the table depicts the relationships between OA isolates. (C) Phylogeny of OA isolates and all Firmicutes assemblies from NCBI GenBank with an ANI 95% to at least one OA isolate. (D-E) UMAP plots of all gene clusters from *Clostridium_AQ innocuum* (D) and *Clostridium perfringens* (E) according to orthogroup presence or absence. OA isolates corresponding to each species are highlighted in red. (F)UMAP plot of KEGG orthologs involved in metabolism for OA14 (in red) and all related *C. perfringens* with an ANI 95%. (G)Number of orthogroups in the core genome (red) and pangenome (teal) of *C. perfringens*, as a function of number of genomes analyzed. Each dot represents the core or pangenome size at a given number of genomes κ, which was computed with up to 100 different combinations of genomes. Heap’s (black lines) was fit to the median core genome (*N* = 6070029 κ^−0.31^) and pan-genome (*N* = 429650κ^0.06^) sizes.

Phylogenetic and taxonomic lineage was identified through whole-genome sequencing and the GTDB-Tk approach for genomic classification (Chaumeil et al., 2019). 13 isolates belonged to Clostridiales, representing five families: *Lachnospiraceae* (OA1, 3, 11, 13), *Peptostreptococcaceae* (OA2, 6, 8), *Erysipelotrichaceae* (OA4, 5, 7), *Oscillospiraceae* (OA9, 10), and *Clostridiaceae* (OA14) (Figure 1B). OA12 was identified as *Enterococcus hirae*. Although *Enterococci* are not spore formers, they can be partially resistant to chemical agents, including chloroform (Maria et al., 2021; Suchomel et al., 2019). All isolates except OA3 had an average nucleotide identity (ANI) of at least 95% to previously sequenced bacteria in public databases, allowing us to assign a species identifier. With an ANI of 88.32% to the closest known relative, OA3 likely represents a new species belonging to the *Ruminococcus* genus, which we have provisionally named *Ruminococcus pangsunibacterium*.

Genome sizes and GC content for OA isolates were generally typical for their taxa and range from 2.85 to 6.95 Mbp and 27.9% to 60.3%, respectively. Phylogenetic comparison to related Firmicutes (Figure 1C) confirmed the taxonomic relation of OA isolates to previously sequenced bacteria. Although some isolates such as OA12 (*Enterococcus hirae*) and OA14 (*Clostridium perfringens*) are extensively studied species, others like OA2 and OA6 (*Romboutsia hominis*) and OA9 and OA10 (genus *Lawsonibacter*) have fewer than 8 other deposited genomes and have yet to be functionally characterized (Figure 1B) (Gerritsen et al., 2018; Sakamoto et al., 2018).

We created UMAP plots of orthogroups of genes for each species to compare the total coding potential of the OA isolates to their relatives (McInnes et al., 2018). We restricted this analysis to species in which a substantial number of sequenced genomes are available: *Clostridium innocuum* (OA4 and OA5), *Parasclostridium sordellii* (OA8), *Enterococcus hirae* (OA12) and *Clostridium perfringens* (OA14). OA4 and OA5 fell into a distinct cluster of *C. innocuum* genomes, while the other OA isolates aligned with the dominant cluster of their respective species (Figure 1D-E, S1B-C). The availability of 622 *Clostridium perfringens* genomes facilitated additional analysis of OA14. A UMAP plot of KEGG orthologs in the metabolism category revealed that OA14 clusters away from most other *Clostridium perfringens* isolates (Figure 1F), suggesting that the metabolic functions of this isolate are distinct.

Quantification of how the pangenome, the whole-genomic repertoire of each microorganism (Vernikos et al., 2015), increases with each added genome revealed that while there have been many genomes sequenced for these species, there is great intra-species diversity and information about these taxa has not yet been saturated (Figure 1G, S1D-G). Thus, OA isolates include both uncharacterized taxa and well-investigated species, which may include those with undefined properties.

### OA genomes are enriched in Malaysian microbiomes

Given that the OA isolates were isolated from two Orang Asli individuals, we tested their broader presence in human microbiomes. We collected stool from a large cohort of 351 Orang Asli from 6 villages, and 56 individuals living in Kuala Lumpur (Tee et al., 2022). The metagenomics classification tool krakenuniq (Breitwieser et al., 2018) was used to map 650 metagenomes generated from the above Malaysian populations to unique sequences from the whole genomes of OA1-14. Whole genomes from the 47 closest relatives of the OA isolates were used to ensure that mapping reflected matches to sequences found exclusively in our isolates (Table S2). We also compared these results with 544 stool metagenomes from the Human Microbiome Project (HMP), which sampled 242 healthy individuals in the United States (Human Microbiome Project Consortium, 2012a; b). We detected higher mapping frequency in the Malaysian metagenomes than HMP metagenomes to sequences specific to OA1-14, as well as to the OA isolates and their close relatives combined (Figures 2A-B). Therefore, the taxa corresponding to the OA genomes are present at higher abundances in the microbiomes of urban and rural Malaysians compared to the microbiomes of Americans.

**Figure 2.**
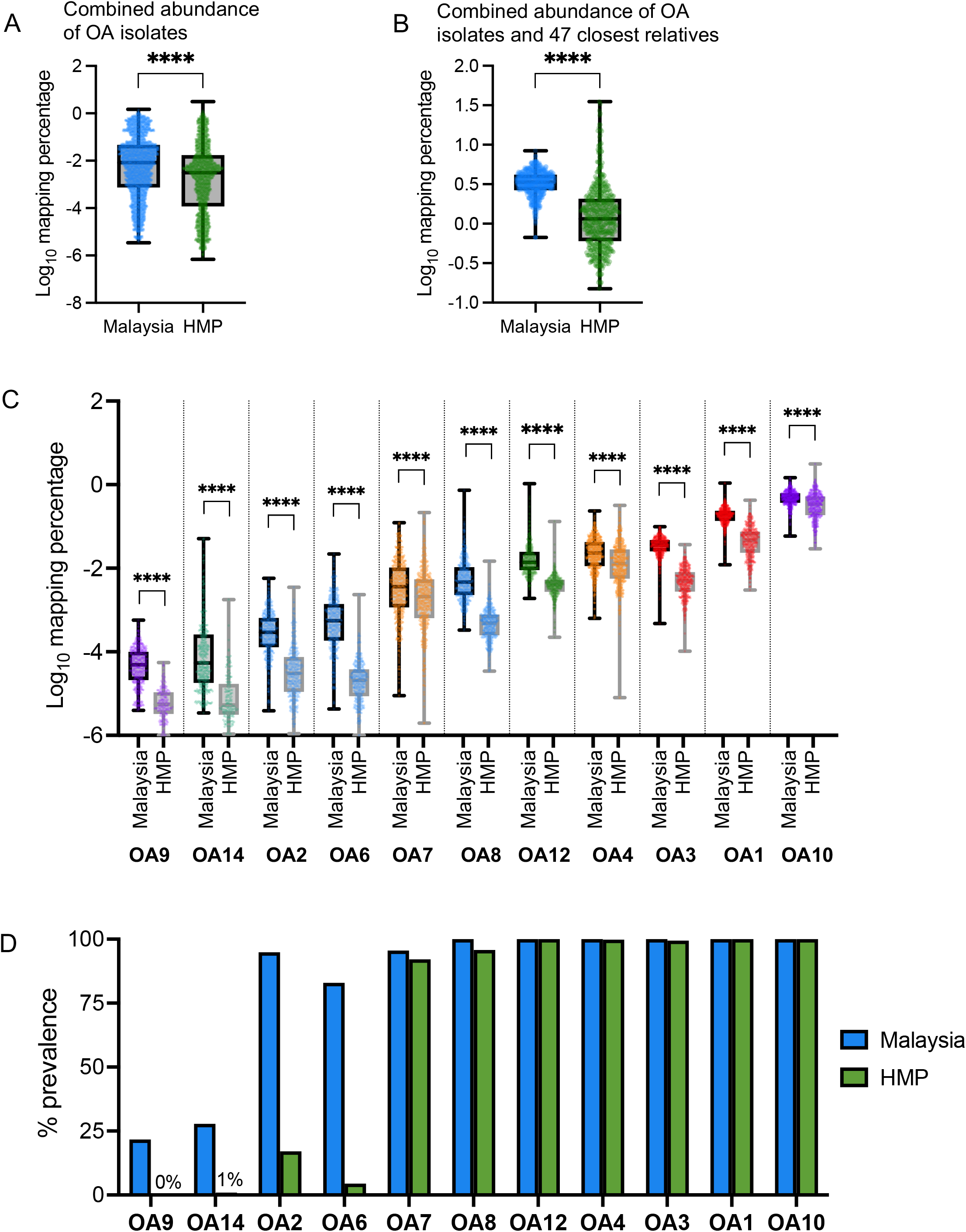
OA genomes are enriched in Malaysian microbiomes. (A) Mapping percentage of Malaysian and HMP metagenomes to genomes of the 14 OA isolates. (B) Mapping percentage of Malaysian and HMP metagenomes to genomes of 14 OA isolates and their 47 closest relatives (see Table S2). (C) Mapping percentage of Malaysian and HMP metagenomes to individual OA isolate genomes. OA5, OA11 and OA13 were not detected by krakenuniq due to low representation. (D) Prevalence of each OA isolate in Malaysian and HMP metagenomes. A mapping percentage of 0.0001% was used as the cutoff for presence of that isolate in each metagenome. (A-C) Each dot represents one metagenome from one individual. Whiskers represent Min and Max. Mapping percentage denotes the percentage of k-mers from each metagenome that specifically map to one of the genomes in the group being measured. Mann-Whitney test, ****p < 0.0001.

Comparing the percentage of Malaysian and HMP metagenomic reads mapped to each individual OA isolate revealed a wide range of mapping percentages indicating different levels of abundance for these taxa. Yet, each isolate displayed significantly reduced abundance in HMP compared with Malaysian microbiomes (Figure 2C). We then analyzed the proportion of microbiomes (prevalence) harboring OA isolates. Seven isolates were equally prevalent (near 100%) in HMP and Malaysian populations, indicating they are ubiquitous microbiome members despite different relative abundance when comparing across individuals in distant geographic regions. In contrast, OA2 and OA6 were rare in HMP metagenomes while displaying high prevalence in Malaysian metagenomes. The two isolates displaying the lowest prevalence in Malaysian metagenomes, OA9 and OA14, displayed very low prevalence or were undetectable in the HMP metagenomes (Figure 2D).

OA11 and OA13 were not detected due to low representation across all metagenomes. As krakenuniq functions by mapping metagenomes to unique sequences in bacterial genomes not found in any other genomes, we hypothesized that OA5 was not detected because its genome is highly similar to OA4. Indeed, when OA4 was removed from the mapping analysis, the mapping percentage of OA5 increased (Figure S2A), indicating that krakenuniq could not differentiate between OA4 and OA5. We detected several positive correlations between pairs of OA isolates within the Malaysian metagenomes, with a particularly strong association between OA2 and OA6, both *Romboutsia hominis* (Figure S2B). The lack of strong negative correlations between isolates suggests these bacteria are not in an antagonistic relationship with each other in this population.

### Identification of OA isolates associated with helminth colonization

The above Malaysian dataset includes individuals living in rural Orang Asli villages and urbanized Kuala Lumpur. The Orang Asli samples can be further segregated into those taken prior to and after treatment with the anthelmintic drug albendazole (ABZ). The helminth prevalence in the Orang Asli population decreased from 67.2% at baseline to 22.5% 21 days after treatment. The helminth prevalence in the urban control population was 0% (Tee *et al.*, 2022). To identify helminth associated bacteria, we determined which OA isolates were enriched in pre-ABZ samples compared with post-treatment and urban control samples. A heat map showing the normalized percentage of k-mers in each metagenome that specifically map to each OA genome revealed that OA2, 6, 8, 12, and 14 were generally rare but exhibited high mapping in a small proportion of individuals, whereas OA1, 3, and 10 had a wider distribution of mapping across individuals (Figure S3A). While OA2, 6, 8, 9, 12 and 14 were more abundant in pre-ABZ Orang Asli villagers than urban controls living in Kuala Lumpur, OA1, 3, 4, 7, 10, and 14 displayed similar or lower abundance in the Orang Asli compared with urban controls, indicating these taxa are associated with the geographical region rather than helminths (Figures 3A-H, S3C-E).

**Figure 3.**
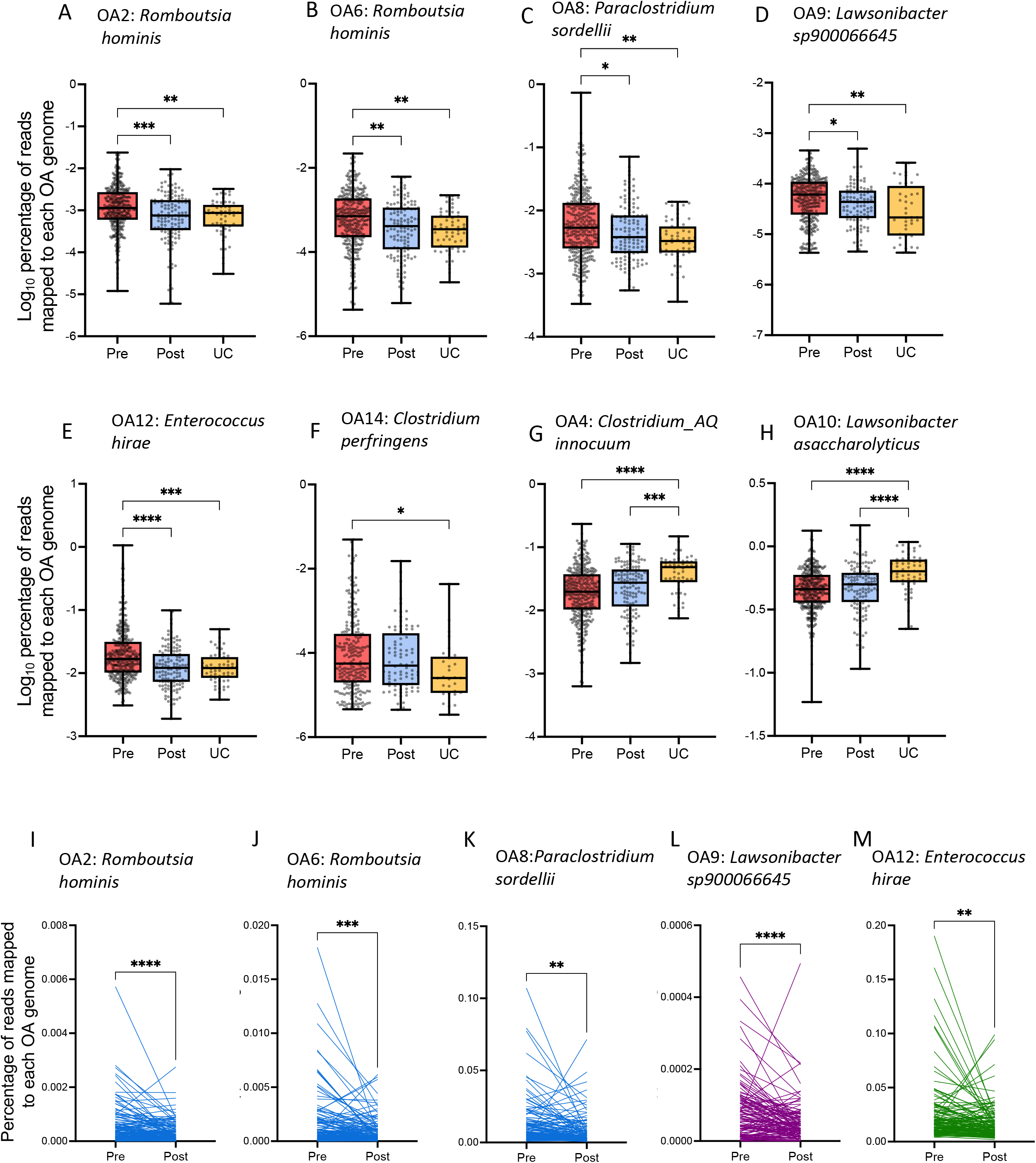
Identification of OA isolates associated with helminth colonization. (A-H) Mapping percentages of pre-albendazole (ABZ), post-ABZ, and urban control (UC) Kuala Lumpur metagenomes to indicated OA isolate genomes. Each dot represents one metagenome from one individual. Whiskers represent Min and Max. (I-M) Pairwise analysis of the mapping percentage of metagenomes derived from matched pre- and post-ABZ longitudinal sampling to indicated OA isolate genomes. Each line represents paired metagenomes from one individual. Kruskal-Wallis with Dunn’s multiple comparisons test used A-H. Paired t-test was used in I-M. *p < 0.05, **p < 0.01, ***p < 0.001, ****p < 0.0001.

Strikingly, all three species belonging to the *Peptostreptococcaceae* family, OA2, OA6, and OA8, had significantly decreased mapping to post-ABZ Orang Asli metagenomes, with mapping percentages to these isolates more closely resembling those in urban controls (Figures 3A-C, S3B). This finding is reminiscent of observations in mice showing that a *Peptostreptococcaceae* species was among the most expanded following *H. polygyrus* inoculation (Rapin et al., 2020). OA9 and OA12 were also reduced in the microbiomes of ABZ-treated individuals (Figures 3D-E, S3B). The remaining OA isolates had no significant change (Figures 3F-H, S3C-E). For 129 individuals, matched pre- and post-ABZ longitudinal samples were available. Pairwise analyses for these samples confirmed that OA2, 6, 8, 9 and 12 were reduced within an individual following ABZ treatment (Figure 3I-M). We also observed decreased OA1 and OA3 abundance within individuals post-ABZ (Figure S3B, F-G), but these isolates were not significantly different between pre-ABZ, post-ABZ, and urban controls at the population level (Figure S3C-D). Considering the relatively high abundance of OA1 and OA3 across all Malaysian metagenomes (Figure 2C), the data suggest these isolates are present at high enough levels to persist in microbiomes in the region despite deworming.

The reduced abundance of OA2, 6, 8, 9, 12 and 14 in the microbiomes of the urban controls from Kuala Lumpur increased our confidence that these taxa are helminth-associated. Although we cannot rule out helminth-independent effects of ABZ, we found that even the highest soluble concentration of ABZ did not impair *in vitro* growth of OA isolates (Figure S3L). Thus, the reduced presence of the above OA isolates in the microbiomes of ABZ-treated individuals is unlikely due to direct killing of these bacteria by the drug.

### Helminth-associated Clostridia induce *Trichuris muris* egg hatching

The host acquires *Trichuris* species following ingestion of embryonated eggs in contaminated food or water, which then hatch to larvae in the microbiota-rich cecum and mature to adult worms. Adult females lay thousands of unembryonated eggs per day that are released through stool to the environment where they embryonate and can transmit to a new host or re-infect the same individual (Bogitsh et al., 2013; Klementowicz et al., 2012). To test the hypothesis that *Trichuris* colonization favors the presence of microbiota members that facilitate its life cycle, we examined whether helminth-associated OA isolates can promote hatching of *T. muris* eggs, the helminth species for which a bacteria-mediated hatching assay is available under aerobic conditions (Hayes *et al.*, 2010; Koyama, 2016). Because Clostridia are obligate anaerobes and the mammalian digestive tract is a low oxygen environment, we first determined whether *T. muris* eggs (Figure 4A) can hatch in the presence of bacteria under anaerobic conditions.

**Figure 4:**
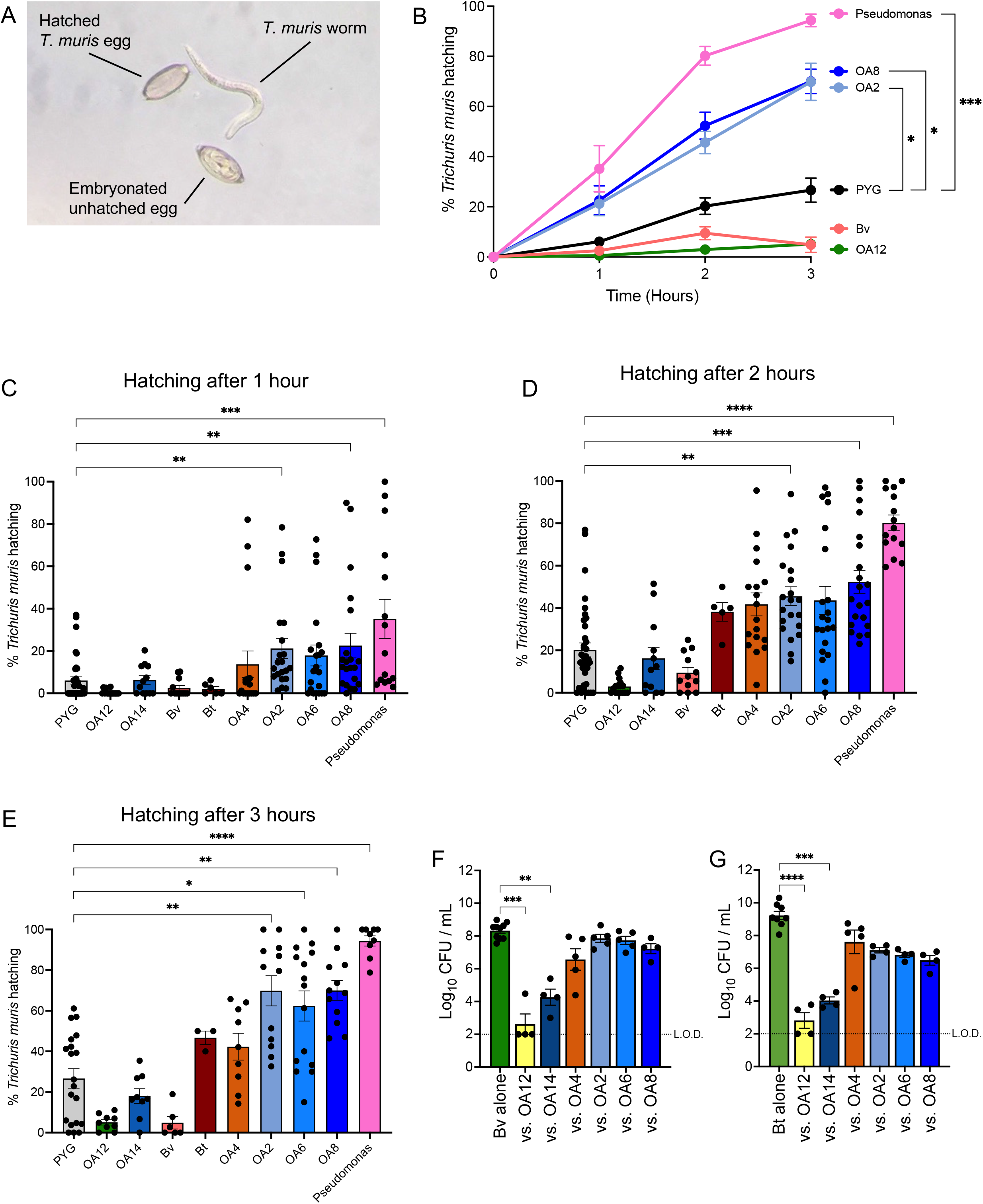
Helminth-associated Clostridia induce *Trichuris muris* egg hatching. (A) Representative light microscopy image of a *T. muris* worm and hatched and unhatched eggs. (B) Time-course of *T. muris* egg hatching in the presence of indicated bacteria. Data is combined from 11 independent experiments, with each condition tested in at least 3 separate experiments, and plotted as mean ± SEM. (C-E) Percentage of hatched *T. muris* eggs after 1 (C), 2 (D) and 3 (E) hours of incubation with indicated bacteria corresponding to the time series graph in (B) to allow visualization of all bacterial strains at each time point. Experiments were performed in at least 3 independent repeats for each strain, and each dot represents one well of eggs. Bars indicate mean ± SEM. (F-G) Colony forming units (CFUs) of *Bacteroides vulgatus* (F) and *Bacteroides thetaiotaomicron* (G) after 24-hours of growth in co-culture with the indicated OA isolates in PYG media. Each dot represents one individual experiment, and bars indicate mean ± SEM. L.O.D., limit of detection = 100 CFU/ml. PYG, Peptone Yeast Glucose media; Pseudomonas, *Pseudomonas aeruginosa;* Bv, *Bacteroides vulgatus*; Bt, *Bacteroides thetaiotaomicron*. Welch’s t-test was used to compare between area under curve of each condition for B. Kruskal-Wallis with Dunn’s multiple comparisons test was used to compare between each OA isolate and PYG in C-E; and between each OA isolate and Bv or Bt in F-G. *p < 0.05, **p < 0.01, ***p < 0.001, ****p < 0.0001.

As a positive control, we chose the facultative anaerobe *Pseudomonas aeruginosa*, which was previously shown to be a potent inducer of hatching in aerobic conditions (Hayes *et al.*, 2010). Incubation of embryonated *T. muris* eggs with 1×10^8^ colony forming units (CFUs) of *P. aeruginosa* resulted in nearly 100% hatching over 3 hours compared with the slower inefficient hatching of <30% for eggs incubated with the peptone yeast glucose (PYG) media alone (Figure 4B). Thus, bacteria can enhance *T. muris* egg hatching under anaerobic conditions. Using this assay, we tested four OA isolates associated with helminths (OA2, 6, 8 and 12) and compared them with two isolates not associated with helminths (OA4 and 14). We also included two Bacteroidales species, *Bacteroides thetaiotaomicron* and *Bacteroides vulgatus*, which are ubiquitous anaerobes in the human microbiome. In addition, *B. vulgatus* growth rate was negatively correlated with *Trichuris* egg burden in the Orang Asli (Tee *et al.*, 2022). We show representative high and low hatching conditions as a time course (Figure 4B) and the full dataset as bar graphs at individual time points (Figure 4C-E) to facilitate visualization. The *Peptostreptococcaceae* strains OA2, 6, and 8 were rapid inducers, with around 20% of eggs hatched after 1 hour of incubation and >60% hatched by 3 hours. Eggs incubated with OA4 and *B. thetaiotaomicron* were slower to hatch, while OA12, OA14, and *B. vulgatus* were poor inducers that potentially inhibit hatching. The degree of hatching did not correlate with the rate of bacterial growth (Figures S4A-C).

Because we previously showed that the increase in Clostridiales in mice and humans colonized with *Trichuris* is associated with a corresponding decrease in Bacteroidales (Ramanan *et al.*, 2016), we tested whether the OA isolates directly impact Bacteroidales growth. We found that OA12 and OA14, two isolates that did not induce *T. muris* egg hatching, were potent inhibitors of *B. vulgatus* and *B. theta* growth when cultured together in an *in vitro* bacterial competition assay (Figures 4F-G). These results indicate that helminth-associated OA isolates, especially *Peptostreptococcaceae* species, promote *T. muris* egg hatching, while competition with *Bacteroides* species may not be a unique property of helminth-associated bacteria.

## Discussion

Whole genome sequencing of the OA isolates and the availability of the largest Malaysian microbiome dataset to date (Tee *et al.*, 2022) allowed us to uncover differential and high-resolution relationships between specific taxa and helminth colonization. Genomes corresponding to OA isolates were enriched in Malaysian metagenomes compared with HMP, yet only a subset of isolates decreased in the microbiomes of individuals treated with deworming medication. The *Peptostreptococcaceae* isolates were particularly notable because they also mediated superior *T. muris* egg hatching. These results raise the possibility that helminth as to be a more hospitable environment for subsequent infections. Individuals living in the same household or environment are known to share microbiome compositions (Song et al., 2013). Thus, naïve hosts may acquire a pro-helminth microbiome by living in proximity with colonized individuals, thereby creating a positive feedback loop that favors propagation of helminths at the community level. A non-mutually exclusive possibility is that eggs shed into the environment are coated by bacteria that promote hatching when ingested together. These hypothetical mechanisms will be important to test in the future as factors that determine helminth burden and susceptibility to reinfection are poorly understood.

Although we validated the presence of the OA isolates in a large metagenome dataset, our focus was limited to a modest number of species. A recent phylogenetic study of *Lachnospiraceae* (family within Clostridiales) isolated from individuals in the U.S. provides compelling evidence of substantial inter- and intra-species diversity in metabolic pathways linked with human health (Sorbara et al., 2020). Given that our OA isolates include understudied taxa and a new species, similar large-scale efforts to isolate and sequence bacteria from the microbiomes of individuals living in low- and middle-income countries are warranted. A second limitation of our study is the use of *T. muris* as a surrogate for the human parasite due to the lack of an established assay to investigate relationships between bacteria and *T. trichiura*. Nevertheless, our observation that bacteria display differential capacity to promote *T. muris* egg hatching provides a key opportunity to investigate mechanisms. Optimal hatching mediated by Proteobacteria under aerobic conditions require fimbriae-dependent attachment to the eggs (Hayes *et al.*, 2010). It will be interesting to determine how Clostridia species, many of which lack fimbriae, promote hatching, and whether the rules are different under anaerobic conditions.

Finally, some of the OA isolates associated with helminths are detected at low levels. Their rarity in the urban controls and HMP microbiomes suggests that transition from a rural lifestyle in a helminth endemic region to an industrialized setting may be leading to a loss of vulnerable low abundance taxa. According to the missing microbes theory, changes in human behavior are leading to loss of beneficial microbes in the gut, contributing to increases in modern diseases (Blaser, 2016; Blaser and Falkow, 2009). Similarly, regions with the highest incidence of immune diseases are associated with the lowest prevalence of helminth infections (Bach, 2017; Elliott and Weinstock, 2012), which may be explained by either direct effects of parasites on the host immune system or indirect effects through the microbiome (Lawson *et al.*, 2021; Loke and Lim, 2015; Maizels, 2020; Maizels et al., 2018; Ramanan *et al.*, 2016). Whether the consequence of losing certain microbes from our microbiome is harmful or reflects an adaptation to modernization remains to be determined (Carmody et al., 2021). Considering the diversity of physiological processes associated with intestinal bacteria, we suggest additional investigation of understudied populations, including those in helminth-endemic regions, will yield valuable knowledge regarding the consequences of a shifting microbiota.

## Supplemental figure legends

**Supplemental Figure 1.**
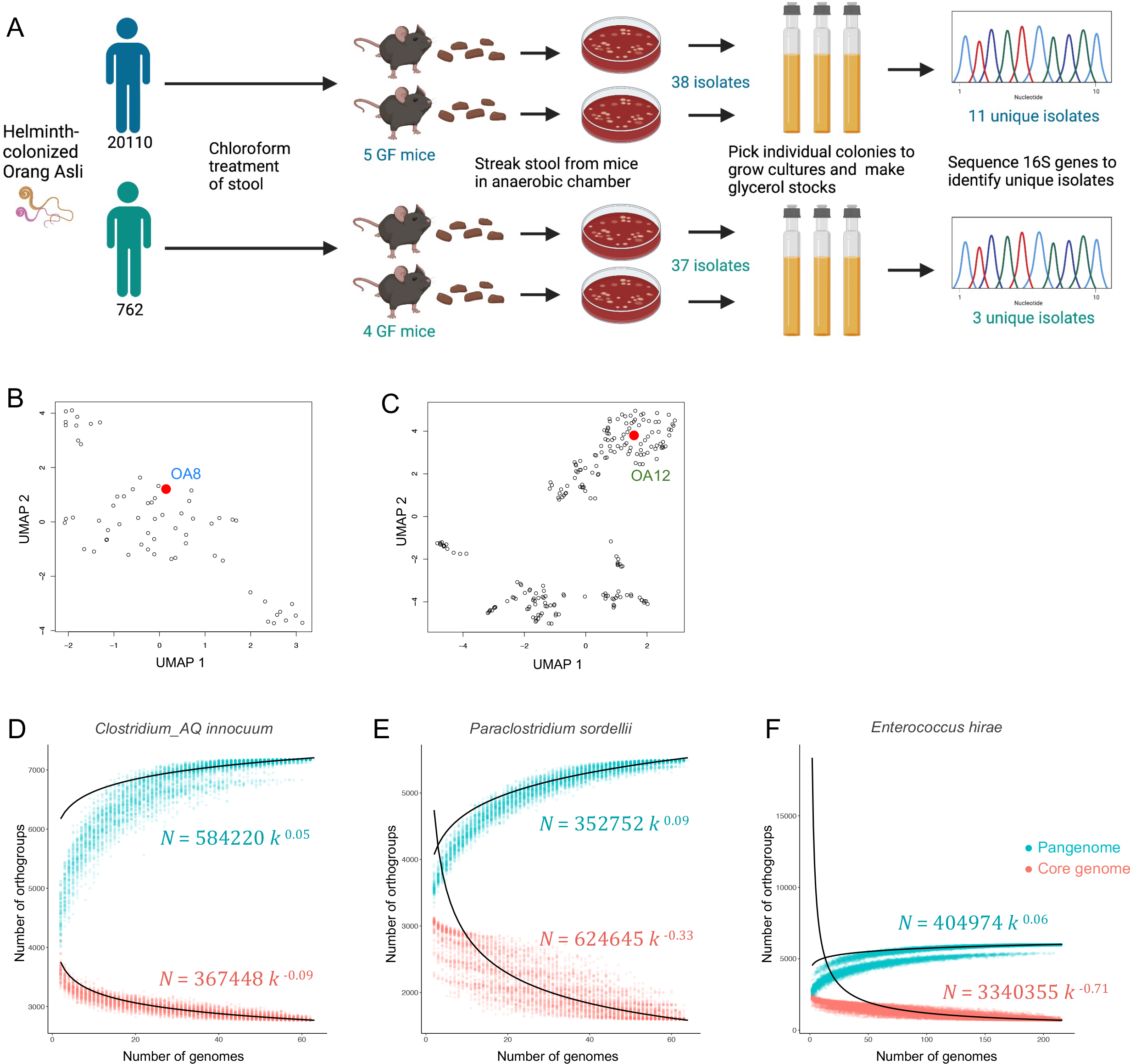
Isolation and identification of spore-forming Firmicutes from helminth-colonized individuals. (A) Schematic for isolation of spore-forming bacteria from the Orang Asli (See Table S1). GF, germ-free. (B-C) UMAP plots of all gene clusters from *Paraclostridium* sordellii (B) and *Enterococcus hirae* (C) according to orthogroup presence or absence. The OA isolate corresponding to each species is highlighted in red. (D-F) Number of orthogroups in the core genome (red) and pangenome (teal) of *Clostridium_AQ innocuum* (D), *Paraclostridium* sordellii (E), and *Enterococcus hirae* (F), as a function of number of genomes analyzed. Each dot represents the core or pangenome size at a given number of genomes κ, which was computed with up to 100 different combinations of genomes. Heap’s law (black lines) was fit to the median core genome and pan-genome sizes; the resulting parameters are indicated on each plot.

**Supplemental Figure 2.**
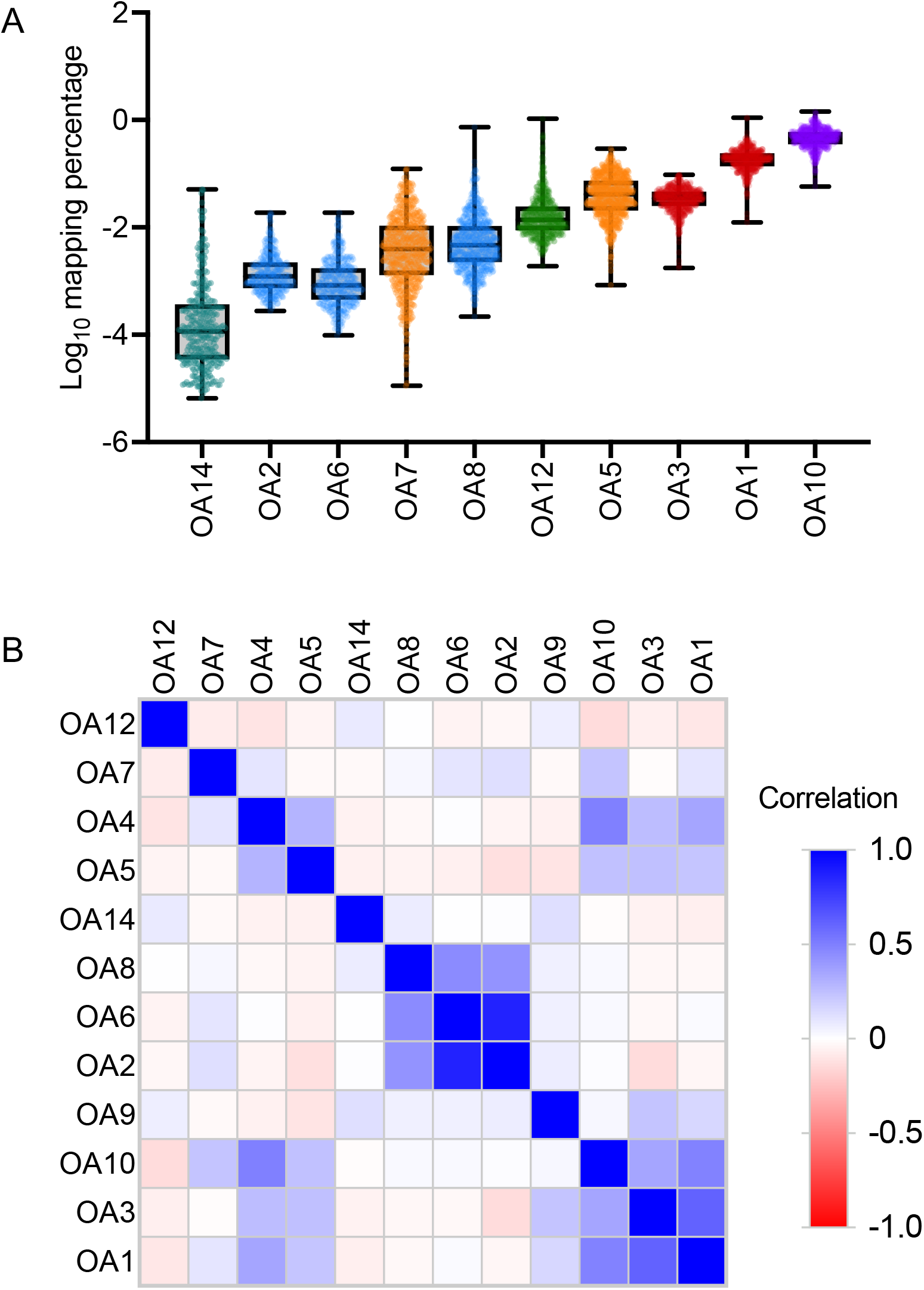
Additional analyses of metagenome mapping to OA genomes. (A) The mapping percentage of OA5 increases when OA4 is removed from the list of genomes during krakenuniq mapping, while mapping percentages of other strains do not change. Each dot represents one metagenome from one individual. Whiskers represent Min and Max. Mapping percentage denotes the percentage of k-mers from each metagenome that specifically map to the genome of each isolate. (B) Heatmap showing Pearson correlations between the mapping percentages of OA isolates in Malaysian metagenomes.

**Supplemental Figure 3.**
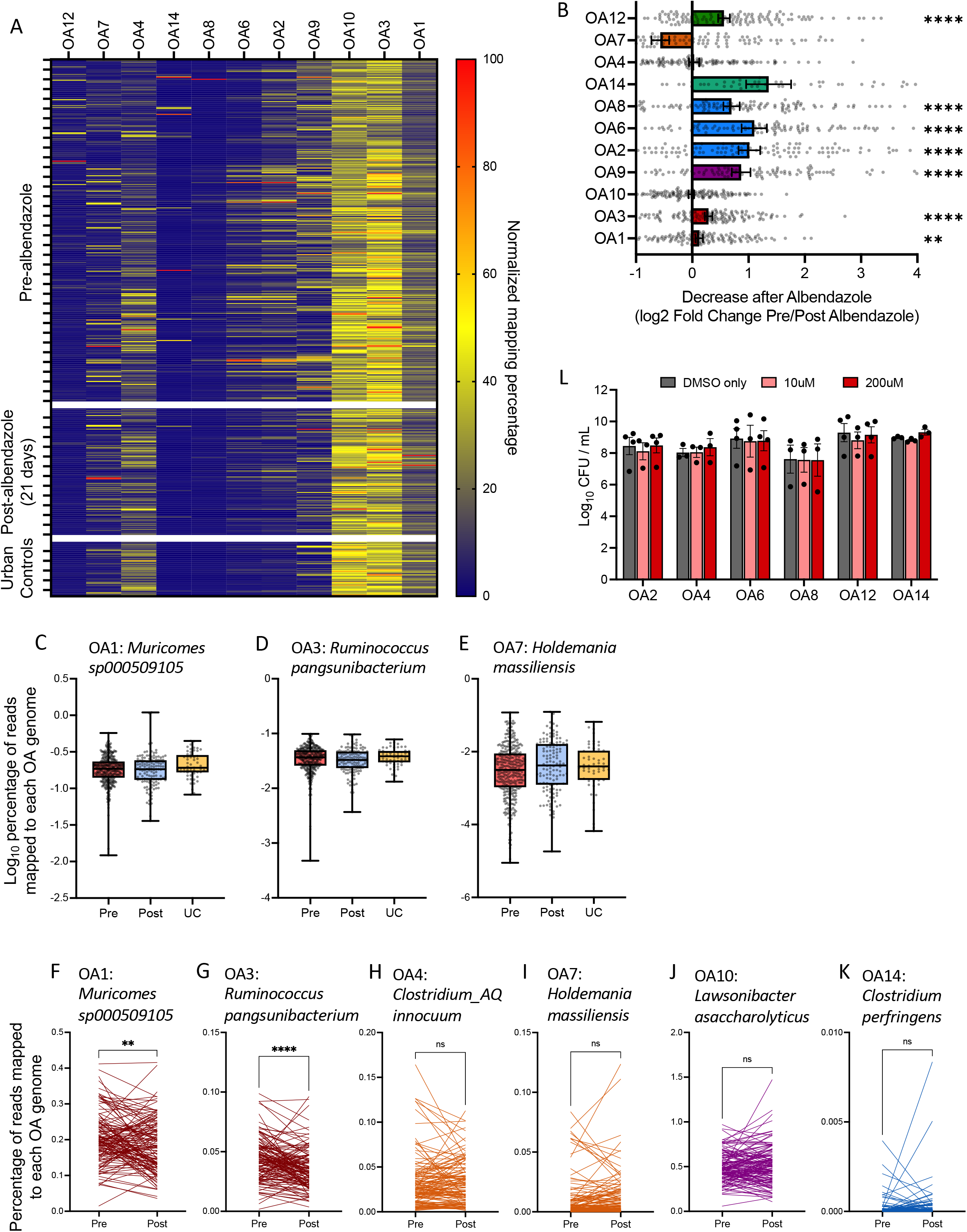
Identification of OA isolates associated with helminth colonization. (A) Heat-map, normalized per column, showing percentage of k-mers in each metagenome that specifically map to each OA genome. (B) Fold-change (pre-ABZ over 21 days post-ABZ) depicting the decrease of the mapping percentage to each OA isolate after ABZ treatment. Each dot represents the fold change in the metagenome of one individual. Error bars represent SEM. (C-E) Mapping percentages of pre-ABZ, post-ABZ, and urban control (UC) Kuala Lumpur metagenomes to indicated OA isolate genomes. Each dot represents one metagenome from one individual. Whiskers represent Min and Max. (F-K) Pairwise analysis of the mapping percentage of metagenomes derived from matched pre- and post-ABZ longitudinal sampling to indicated OA isolate genomes. Each line represents paired metagenomes from one individual. (L) Colony forming units (CFUs) of indicated bacteria grown in the presence of 10uM or 200uM ABZ or DMSO vehicle control. Dots represent averages of individual experiments and error bars represent SEM. (B) One sample Wilcoxon test comparing against a hypothetical value of fold change = 1; **p < 0.01 and ****p < 0.0001 (C-E) Kruskal-Wallis with Dunn’s Multiple comparisons test; no significance. (F-K) Paired t-test; **p < 0.01, ****p < 0.0001. (L) Mixed-effects model with Tukey’s multiple Multiple comparisons test revealed no significant difference between treatment conditions for all isolates.

**Supplemental Figure 4.**
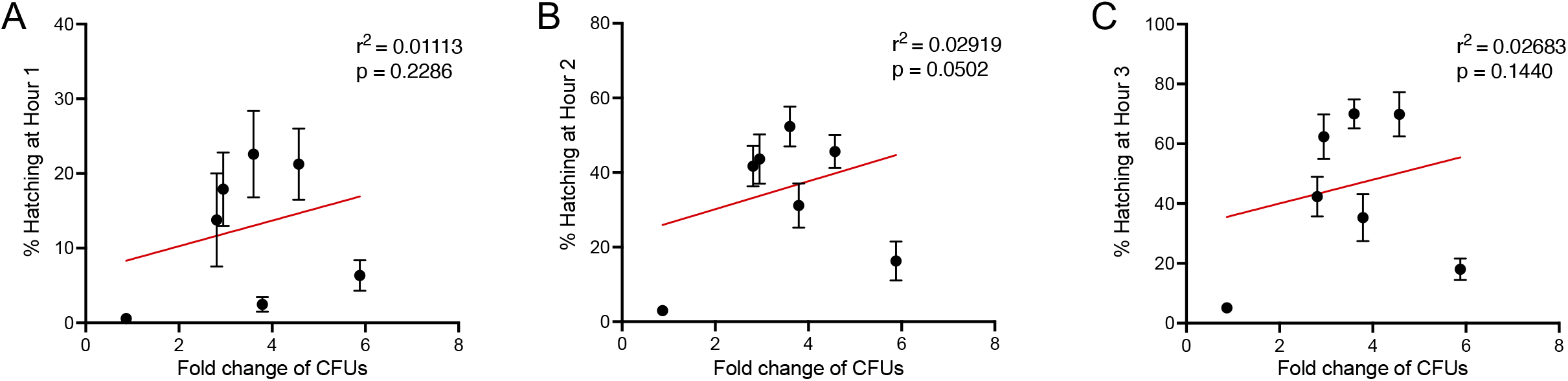
Additional analyses of OA isolates for properties associated with Trichuris muris egg hatching. (A-C) Linear regression analysis comparing the fold change of bacterial growth over 4 hours and the corresponding percentage of hatched *T. muris* eggs by that bacterium after 1 (A), 2 (B) and 3 (C) hours. Each dot represents one bacterial isolate and error bars represent SEM.

## Methods

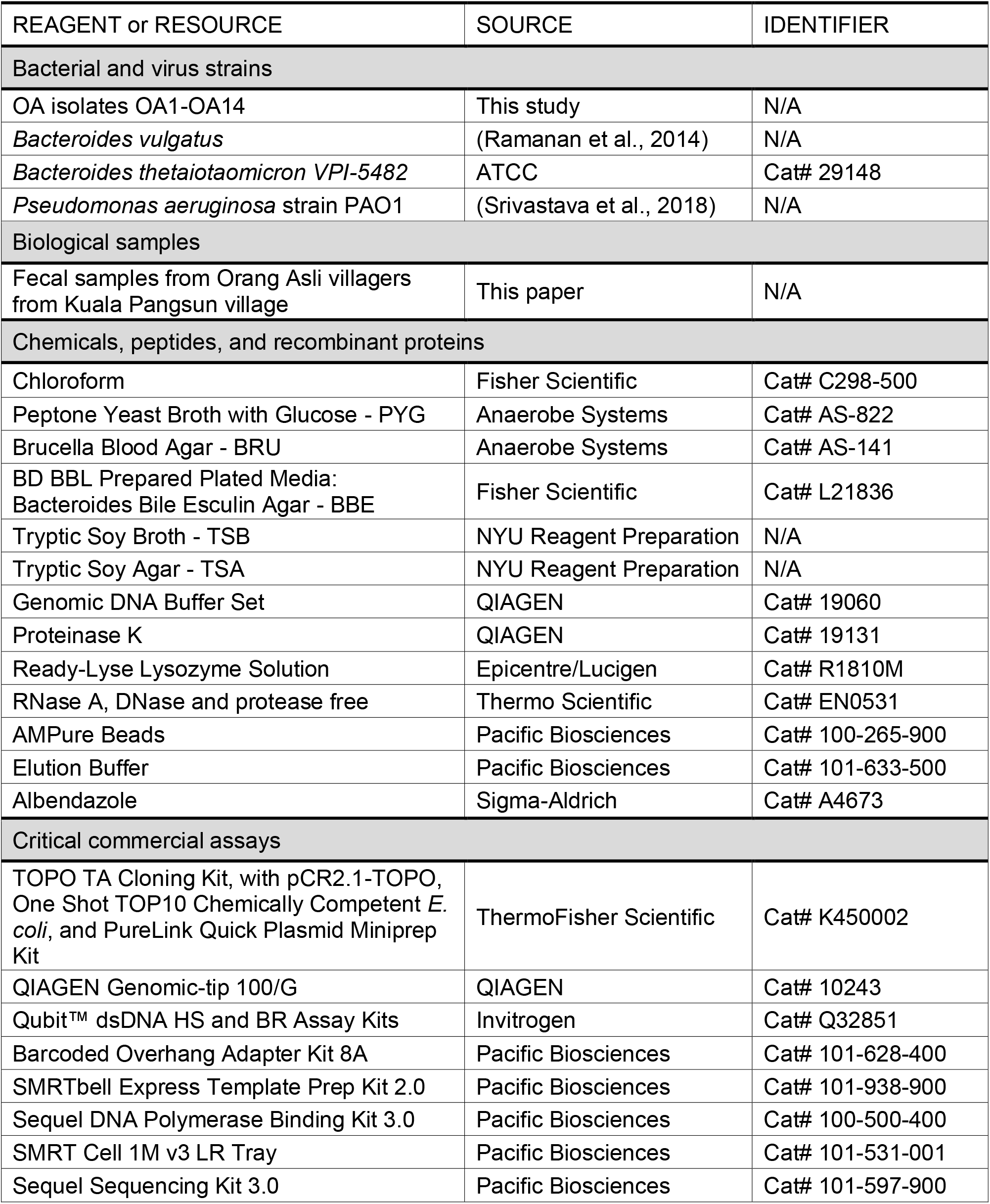
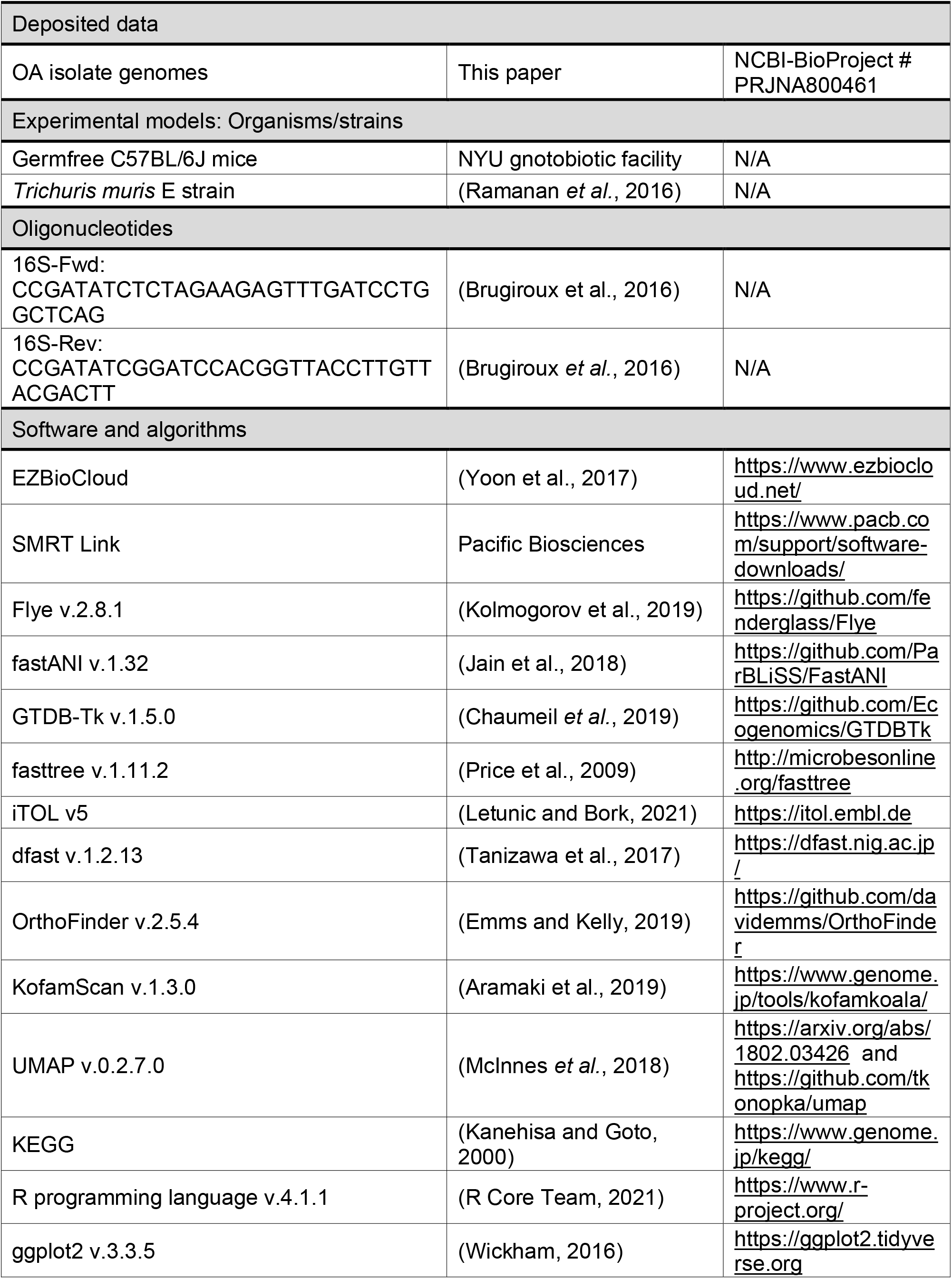
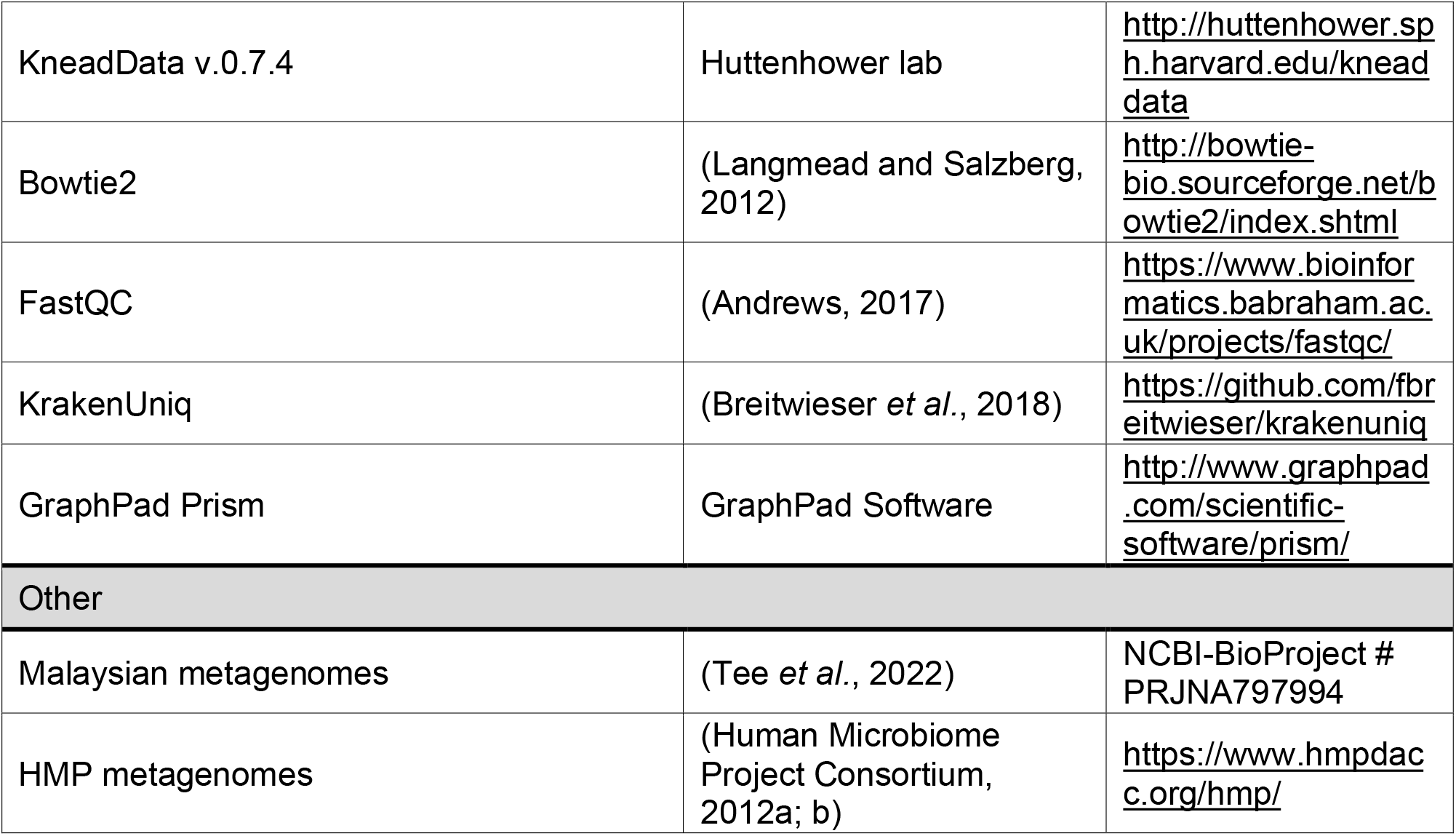
Key resources table.

### Contact for reagent and resource sharing

Further information and requests for resources and reagents should be directed to and will be fulfilled by Lead Contact, Dr. Ken Cadwell. All unique and stable reagents generated in this study are available from the Lead Contact with a completed Materials Transfer Agreement.

## EXPERIMENTAL MODEL AND SUBJECT DETAILS

### Human subjects

Prior to collection of samples from Kuala Pangsun village (Table S1), the human study was approved by the Research and Ethics Committee of Universiti Malaya Medical Center (UMMC) (i.e., MEC Ref. No. 824.11, No. 943.14 and No. 2017925-5593), National Medical Research Register (NMRR), Ministry of Health, Malaysia (Reference No.: NMRR-17-3055-37252), the Department of Orang Asli Development (JAKOA) [Ref. No.: JAKOA/pp.30.052Jld13 (12) & JAKOA/pp.30.052Jld14 (47)], and NYU IRB# i17-01068. Permission was also obtained from the Tok Batin, chieftain of the village. Kuala Pangsun village (101.88°E longitude, 3.21°N latitude) is situated in Hulu Langat district, the fifth largest district in Selangor state, Malaysia. The selection of this village was favorable in terms of logistics and feasibility, coupled with good cooperation from the villagers. The purpose and the procedure of this study was explained orally to all the participants by the investigator. Written consent was attained from all adult participants aged 18 and above. For children under 18 years old, written parental consent was obtained from their respective parents or guardian. Study exclusion criteria consisted of pregnant women, breastfeeding mothers and presence or perceived presence of any clinically significant disease. Information about human subjects from the large Orang Asli and Kuala Lumpur cohort which provided metagenomes for our analyses is described in (Tee *et al.*, 2022).

### Germfree mice

Germfree (GF) C57BL/6J were bred and maintained in flexible-film isolators at the New York University Grossman School of Medicine Gnotobiotics Animal Facility. Absence of fecal bacteria was confirmed monthly by evaluating the presence of 16S DNA in stool samples by qPCR as previously described (Kernbauer et al., 2014). For inoculation with spore-forming stool fractions, GF mice were housed in Bioexclusion cages (Tecniplast) with access to sterile food and water. All animal studies were performed according to protocols approved by the NYU Grossman School of Medicine Institutional Animal Care and Use Committee.

### Bacterial strains

In addition to the OA isolates from this study, *Bacteroides vulgatus* was isolated by our lab previously (Ramanan *et al.*, 2014), *Bacteroides thetaiotaomicron* (VPI-5482) was kindly provided by E. Martens (University of Michigan Medical School), and *Pseudomonas aeruginosa* was kindly provided by A. Darwin (NYU Grossman School of Medicine) (Srivastava *et al.*, 2018). All bacteria except *P. aeruginosa* were cultured under anaerobic conditions in an anaerobic chamber (Coy Labs). Frozen glycerol stocks (30% glycerol) of all bacteria were prepared. Glycerol stocks of the OA isolates and *Bacteroides* species were streaked onto BRU plates (Anaerobe Systems) and incubated anaerobically for 48 hours at 37°C. PYG broth (Anaerobe Systems) inoculated with single colonies was grown at 37°C. OA1, 2, 4, 5, 6, 8, 11, 12, 13, and 14 were grown for 24 hours. OA3, 7, 9, and 10 required 3 days to reach similar turbidity. *P. aeruginosa* glycerol stocks were streaked onto TSA plates and incubated aerobically for 24 hours at 37°C, after which single colonies were spiked into TSA broth and grown for 24 hours at 37°C with shaking at 225 rpm. To quantify colony forming units, we performed serial dilutions of liquid culture in sterile PBS and plated on BRU agar for the OA isolates, BBE agar (BD) for *Bacteroides* species, and TSA agar for *P. aeruginosa.*

### Parasite maintenance

Stock eggs of *Trichuris muris* E strain (Ramanan *et al.*, 2016) were propagated and maintained as previously described (Antignano et al., 2011). Each egg batch was confirmed to hatch at ≥80% *in vitro* using the method described in (Venzon *et al.*, 2021) before use in subsequent experiments.

## METHOD DETAILS

### Sample collection from Orang Asli

A total number of 19 fecal samples were collected from 19 individuals (9 female, 10 male, ages 5-51) (Table S1). Participants were given screw-capped containers labeled with names. Containers with participant’ samples were collected the following day, immediately frozen on dry ice, and transported on dry ice to the Department of Parasitology, Faculty of Medicine, Universiti Malaya. Fecal samples were then separated into three portions: (i) preserved in 2.5% potassium dichromate and stored at 4°C for intestinal helminth infection screening, (ii) aliquoted into anaerobic glass hungate tubes for isolation of bacteria, and (iii) aliquoted in 1.5ml cryovial tubes for long-term storage of samples. Aliquots (ii) and (iii) were transported on dry ice to NYU Grossman School of Medicine while aliquot (i) was kept at the University of Malaya to quantify helminth burden using Kato-Katz and formalin ether as described in (Ramanan *et al.*, 2016)

Sample collection and processing for the large Orang Asli and Kuala Lumpur cohort which provided metagenomes for our analyses is described in (Tee *et al.*, 2022). In brief, fecal samples were collected from Orang Asli villagers right before administration of albendazole (ABZ) at 400mg/day for 3 consecutive days and follow-up samples were collected after 21 days. Individuals in urban Kuala Lumpur were not given ABZ and samples were only collected at one timepoint.

### Isolation and identification of OA isolates from human stool

Spores were enriched from stool using a previously published protocol (Atarashi *et al.*, 2013). Fecal samples frozen in hungate tubes were moved into an anaerobic chamber (Coy Labs). ~50mg of the sample was suspended in 10 times volume (w/v) of sterile pre-reduced PBS and passed through a 70 mm cell strainer. Suspensions were mixed with chloroform at a final concentration of 3%, shaken vigorously for 30 seconds, then incubated at 37°C for 1 hour. Chloroform was then removed through evaporation by bubbling with N_2_ gas for 30 min. Suspensions from each donor were inoculated into 4-5 germ-free C57BL/6J mice by intra-gastric gavage (250ul per mouse). Stools on day 7 post-inoculation were collected and suspended in sterile PBS, homogenized in a bead beater in safe-lock tubes with 1.0 mm silica beads, streaked onto BRU agar in an anaerobic chamber, and incubated for 2-4 days at 37°C. 75 colonies were selected, with the aim of picking as many different morphologies as possible, spiked into PYG broth, incubated for 1-3 days at 37°C, and used to prepare sterile glycerol stocks. The inoculation loop used to pick each isolate was dipped into PCR grade water to perform a 16S colony PCR on each isolate using primers 16S-Fwd and 16S-Rev (Brugiroux *et al.*, 2016) to remove duplicates from the pool. After identifying 14 isolates with unique 16S sequences and/or morphologies, we extracted DNA from liquid cultures of these isolates using phenol-chloroform, then subcloned and sequenced the full 16S gene using the primers 16S-Fwd and 16S-Rev and the TOPO™ TA Cloning™ kit with pCR™ 2.1-TOPO™ plasmid (ThermoFisher). Sequences were aligned to the EzBioCloud 16S rRNA database (Yoon *et al.*, 2017) for initial taxonomic identification.

### Whole-genome sequencing and genome assembly

DNA was isolated from liquid culture of all isolates and lysed overnight in Buffer B1 (QIAGEN) containing 200ug/ml RNase A (ThermoScientific), 1250U/ul lysozyme (Epicentre) and 100ul Proteinase K (QIAGEN), at 37°C. High molecular weight DNA was isolated using the Genomic-tip 100/G kit (QIAGEN) and run on the Agilent 2200 TapeStation system with Genomic ScreenTape (cat# 5067-5365) to verify large fragment DNA. Once verified, 1.5ug of DNA was sheared using the Covaris G-TUBE (cat# 520079) and spun for 2min in each direction at 5.4rpm in the Eppendorf 5415 D centrifuge to shear the DNA to an average size of 13,000bp length. Each sheared sample was barcoded with the PacBio Barcoded Overhang Adapter Kit 8A (cat# 101-628-400) during library prep using the PacBio SMRTbell Express Template Prep Kit 2.0 (cat# 100-938-900). After barcoding, each sample was verified on the Invitrogen Qubit system with the dsDNA HS kit (cat# Q32854) for concentration and on the Agilent 2200 TapeStation system for size. Samples were combined to an equal concentration using the PacBio Express microbial multiplexing calculator to give an estimated size of about 30mb total. The PacBio Sequencing Primer v4 was conditioned and annealed to the pooled sample based on the Diffusion loading instructions found on the PacBio SMRT Link software version 8.0.0805529. Following primer binding, the diluted PacBio Sequel Polymerase 3.0 (cat# 100-500-400) was bound to the sample. After binding, the sample was cleaned with PacBio Ampure Beads (cat# 100-265-900) and eluted in PacBio Elution Buffer (cat# 101633-500), and the final concentration of the pool was verified on the Invitrogen Qubit system as before. The pool was combined with final loading reagents (part of the Express kit) and run on CLR mode (continuous long reads) on the PacBio Sequel using a 20hr run time with the SMRT Cell 1M v3 LR tray (cat# 101-531-001) and the Sequel Sequencing Kit 3.0 reagent plate (cat# 101-597-900). Following the sequencing, all genomes were assembled through SMRT Link software except OA2. The OA2 genome was assembled using Flye 2.8.1 in pacbio-raw mode with asm-coverage 150, sub-sampling the reads for the initial assembly to a coverage of 150x (Kolmogorov *et al.*, 2019).

### Species identification, phylogenetic analysis, orthogroup and KEGG UMAP plots, and pangenome plots

To obtain a phylogenetic context for the OA isolates, a total of 190,173 genome assemblies that had been assigned to the phylum Firmicutes (NCBI taxonomy ID: 1239) were downloaded from NCBI GenBank (https://www.ncbi.nlm.nih.gov/genbank/) on November 24, 2021. Of those, only 1,009 were retained after comparing them to all assemblies of the OA isolates with fastANI v.1.32 (Jain *et al.*, 2018) and filtering for estimated average nucleotide identity (ANI) values ≥0.95. The workflow classify_wf from GTDB-Tk v.1.5.0 (Chaumeil *et al.*, 2019) was applied to 1,023 retained GenBank and OA assemblies to obtain a taxonomic classification, using default parameters. We adopted GTDB-Tk taxonomic designations. Using fasttree v.1.11.2 (Price *et al.*, 2009), a phylogenetic tree was calculated with the alignment of concatenated marker genes that was produced by GTDB-Tk from all 1,023 assemblies, using default parameters. Phylogenies were plotted with iTOL (Letunic and Bork, 2021).

The 1,023 assemblies (above) were annotated with dfast v.1.2.13 (Tanizawa *et al.*, 2017) and grouped by GTDB-Tk species designation. Orthofinder v.2.5.4 (Emms and Kelly, 2019) was used with default parameters for computing orthogroups (also called clusters of orthologous genes) for each individual species group and for all 14 OA assemblies as a group. The phylogenetic tree in Figure 1B was produced by orthofinder using the group of OA assemblies only as an input. However, the largest group of 623 assemblies classified as *Clostridium_P perfringens* could not be processed with orthofinder due to its size. For this reason, a subset of 221 assemblies was chosen from the 623 such that the ANI between the assemblies in the subset was maximized. KEGG orthologs were mapped to the predicted proteomes of each assembly (dfast, above) using KofamScan v.1.3.0 and HMM profiles downloaded from KEGG on June 24, 2021 (Aramaki *et al.*, 2019; Kanehisa and Goto, 2000). The resulting sets of KEGG ortholog identifiers were mapped and annotated with all KEGG BRITE hierarchies released on March 30, 2020.

To produce pangenome plots for each species, the numbers of orthogroups in the core and pan-genome were successively computed for up to 100 size-*k* subsets of the available assemblies. Specifically, we started with κ = 2 and successively increased until it reached the number of available assemblies for a species, *n*. For each κ, the number of size-κ combinations of genomes, *n*Cκ was computed. If *n*Cκ ≤ 100, all possible cobination times of the assemblies were considered when computing core and pan-genome sizes. If *n*Cκ > 100, long computation times and combinatorial explosion were prevented by only considering 100 size-random samples of the assemblies that were drawn without replacement. Using linear regression, two Heap’s law curves *N* = *β*κ^*α*^ were fit for each species: one for the median core genome sizes and one for the median pangenome sizes. Only 50% highest values of k were used for fitting Heap’s law curves (Tettelin et al., 2008).

The R programming language v.4.1.1 (R Core Team, 2021) was used for plotting, using ggplot2 v.3.3.5 (Wickham, 2016). UMAP dimensionality reduction was carried out using UMAP v.0.2.7 within R (https://github.com/tkonopka/umap). For the orthogroup plots of each species, the input to UMAP was a binary matrix indicating presence or absence of an orthogroup in each strain. For the plot of KEGG metabolic genes in *Clostridium_P perfringens*, the number of KEGG orthologs in the KEGG brite hierarchy *09100 Metabolism* were tabulated according to the categories of the most specific level of this hierarchy, *D*, and a matrix indicating the numbers of genes in each category and strain was input to UMAP.

### Metagenome classification using Krakenuniq

Malaysian metagenomes were obtained from NCBI-BioProject # PRJNA797994 (Tee *et al.*, 2022) and metagenomes from the Human Microbiome Project (HMP) were obtained from https://www.hmpdacc.org/hmp/ (Human Microbiome Project Consortium, 2012a; b). The file type used in the HMP data portal was wgs_raw_seq_set and we only used fecal samples from healthy human subjects. Quality control, trimming and human host genome removal were performed by KneadData v.0.7.4 with default settings. In KneadData, sequence reads were mapped against human reference genome (hg37) using Bowtie2 with default parameters (very-sensitive end-to-end alignment) to remove the human host genome (Bolger et al., 2014; Langmead and Salzberg, 2012). Human host genome removal was not performed on HMP metagenomes because the human reference genome had already been filtered in the HMP data. Next, FastQC was used to verify quality control (Andrews, 2017). Krakenuniq was used for taxonomic classification (Breitwieser *et al.*, 2018). The cleaned reads from the Malaysia dataset and HMP were mapped against the OA isolate genomes and their 47 closest relatives (Table S2). In Figure S2A, classifications with lower than 50 k-mer counts were removed due to low confidence.

### *T. muris* egg hatching assay

25ul embryonated *T. muris* eggs at a concentration of 1 egg/1ul in sterile water were mixed with 15ul sterile PYG media and 10ul overnight bacterial culture in triplicates in a 48 well plate. PYG-control wells contained an additional 10ul sterile PYG media instead of bacterial culture. Plates were incubated at 37°C in an anaerobic chamber and hatched eggs were quantified every hour over the course of three hours on the Zeiss Primovert microscope, by counting hatched and embryonated unhatched eggs in each well. Unembryonated eggs, which lack visible larvae and have disordered contents, were excluded due to their inability to hatch. A separate 48-well plate was used per timepoint as the plate needed to be removed from the anaerobic chamber to count colonies at each timepoint.

### Albendazole assay

Albendazole (ABZ) (Sigma) was dissolved in DMSO to make 2.5mM and 500mM stocks that were diluted with sterile PYG to a final concentration of 10uM and 200uM. Concentrations used in *in vitro* experiments were chosen based on max solubility in DMSO and max amount of DMSO added to culture (0.5%). Single colonies of bacteria were spiked into 5ml PYG containing 10um ABZ, 200uM ABZ or 0.5% DMSO as a vehicle control, in duplicates in sterile polypropylene tubes. After incubation for 24 hours at 37°C in anaerobic conditions, serial dilutions of the bacterial culture in sterile PBS were plated on BRU agar for OA isolates and BBE agar for *Bacteroides* species and incubated for 48 hours to count colonies and calculate CFUs/ml of culture.

### Bacterial competition assay

Single colonies of *B. vulgatus* or *B. thetaiotaomicron* were inoculated into duplicates of 7ul PYG media along with single colonies of OA isolates. Cultures were grown for 24 hours at 37°C under anaerobic conditions after which serial dilutions of the bacterial culture in sterile PBS were plated on BRU agar for Clostridia and BBE agar for Bacteroides and incubated for 48 hours to count colonies and calculate CFUs/ml of culture.

## QUANTIFICATION AND STATISTICAL ANALYSIS

For *in vitro* experiments, the number of repeats per group is annotated in corresponding figure legends. Significance for all experiments was assessed using GraphPad Prism software (GraphPad). Specific tests are annotated in corresponding figure legends. P values correlate with symbols: ns or no symbol = not significant, *p < 0.05, **p < 0.01, ***p < 0.001, ****p < 0.0001.

## DATA AND SOFTWARE AVAILABILITY

PacBio raw sequences and assembled whole genome sequences reported in this study are available under NCBI-BioProject # PRJNA800461.

## Acknowledgements

We would like to thank Margie Alva, Juan Carrasquillo and David Basnight for their help in the NYU Gnotobiotic Facility, Paul Zappile, Dacia Dimartino, Kim Castelli and Christian Marier at the NYU Langone Genome Technology Center for their help with whole genome sequencing and assemblies, and the NYU Reagent Preparation service for providing bacterial media. We thank Gregory Putzel for help with genome analyses, Menghan Liu for help with krakenuniq, and Mericien Venzon for providing *T. muris* eggs. We would also like to thank Juan Lafaille and members of the Cadwell and Loke Labs for their constructive comments. Figure S1A was created using BioRender.com. This work was in part funded by NIH grants DK093668 (K.C.), HL123340 (K.C.), AI130945 (K.C., P.L., Y.L.), AI140754 (K.C.), DK124336 (K.C.), AI121244 (K.C.), and AI133977 (P.L., V.J.T.). Further funding was provided by Faculty Scholar grant from the Howard H M a I (K.C.), C & C F a (K.C.), Kenneth Rainin Foundation (K.C.), Judith & Stewart Colton Center of Autoimmunity (K.C.), NIH grant 2T32AI007180 (S.S.), and the NYU Langone Health Antimicrobial-Resistant Pathogens Program (A.P.). PacBio sequencing reagents were partially paid for through the PacBio Local SMRT Grant (K.C.). NYU La G T C a M La a are supported by NYU Cancer Center support grant P30CA016087 and further support for the PacBio Sequel was provided by NIH Shared Instrumentation Grant 1S10OD023423-01.

## Author contributions

S.S., P.L. and K.C. conceived and designed the study. M.Z.T., Y.X.E. and Y.A.L.L. carried out and oversaw the collection of samples from Malaysia. S.S. performed the experiments and analyzed and interpreted the data. A.R. and J.S. helped perform *T. muris* hatching experiments. A.H. oversaw whole genome sequencing. A.P. and R.C. assembled and analyzed bacterial genomes. Z.C. and J.C.D. ran the metagenomic mapping analysis with guidance from K.V.R. and V.J.T. S.C.L. provided metadata and help with Malaysian metagenome datasets. K.C., P.L. and S.C.L. oversaw analysis and interpretation of all experiments described. S.S. and K.C. wrote the paper with input from all authors.

---

**Table S1.** Sample characteristics of the Orang Asli cohort from Pangsun village from which bacteria were isolated. ‘*Trichuris* Formalin either’ indicates whether a worm was detected in each stool sample (1) or not (0) and *Trichuris* Kato Katz Intensity’ indicates the number of helminth eggs per gram in each stool sample. The OA strains in this study were isolated from individuals #20110 and #762 (underlined).

| **Sample ID** | **Age** | **Gender** | ***Trichuris* Formalin ether** | ***Trichuris* Kato Katz Intensity** |
| --- | --- | --- | --- | --- |
| <u>20110</u> | <u>5</u> | <u>M</u> | <u>1</u> | <u>4375</u> |
| 88 | 9 | M | 1 | 3850 |
| 135 | 12 | M | 1 | 2450 |
| <u>762</u> | <u>9</u> | <u>F</u> | <u>1</u> | <u>420</u> |
| 1097 | 12 | F | 1 | 385 |
| 37 | 8 | M | 1 | 315 |
| 4 | 5 | M | 1 | 315 |
| 19100 | 11 | M | 1 | 175 |
| 794 | 6 | F | 1 | 140 |
| 1999 | 6 | F | 1 | 35 |
| 445 | 12 | M | 0 | 0 |
| 1889 | 36 | F | 0 | 0 |
| 930 | 16 | F | 0 | 0 |
| 218 | 51 | M | 0 | 0 |
| 995 | 9 | F | 0 | 0 |
| 1612 | 33 | F | 0 | 0 |
| 119 | 50 | M | 0 | 0 |
| 98 | 35 | F | 0 | NA |
| 1836 | 7 | M | 0 | NA |

**Table S2.** 47 bacterial genomes with highest sequence similarity to the OA isolates OA1-14, used in krakenuniq metagenomic mapping to ensure specificity of sequence matches to OA1-14.

|  |  |
| --- | --- |
| 1. | Anaerobutyricum_hallii_isolate_EH1.fna |
| 2. | Clostridioides_difficile_CD196.fna |
| 3. | Clostridioides_difficile_strain_DSM_28668.fna |
| 4. | Clostridium_acetobutylicum_ATCC_824.fna |
| 5. | Clostridium_argentinense_strain_89G.fna |
| 6. | Clostridium_beijerinckii_ATCC_35702_strain_SA-1.fna |
| 7. | Clostridium_botulinum_A_str._ATCC_3502.fna |
| 8. | Clostridium_botulinum_A_str._Hall.fna |
| 9. | Clostridium_botulinum_B1_str._Okra.fna |
| 10. | Clostridium_botulinum_F_str._Langeland.fna |
| 11. | Clostridium_botulinum_strain_Mfbjulcb3.fna |
| 12. | Clostridium_botulinum_strain_Mfbjulcb6.fna |
| 13. | Clostridium_botulinum_strain_NCTC_8550.fna |
| 14. | Clostridium_butyricum_60E.3_actYk-supercont1.1.fna |
| 15. | Clostridium_carboxidivorans_P7.fna |
| 16. | Clostridium_chauvoei_strain_DSM_7528.fna |
| 17. | Clostridium_drakei_strain_SL1.fna |
| 18. | Clostridium_formicaceticum_strain_ATCC_27076.fna |
| 19. | Clostridium_formicaceticum_strain_DSM_92.fna |
| 20. | Clostridium_kluyveri_DSM_555.fna |
| 21. | Clostridium_kluyveri_NBRC_12016.fna |
| 22. | Clostridium_ljungdahlii_DSM_13528.fna |
| 23. | Clostridium_pasteurianum_DSM_525_=_ATCC_6013.fna |
| 24. | Clostridium_perfringens_str._13.fna |
| 25. | [Clostridium]_saccharolyticum_WM1.fna |
| 26. | Clostridium_sp._D5_supercont1.1.fna |
| 27. | Clostridium_sp._SY8519.fna |
| 28. | Clostridium_tetani_E88.fna |
| 29. | Clostridium_tyrobutyricum_strain_KCTC_5387.fna |
| 30. | Escherichia_coli_str._K-12_substr._MG1655.fna |
| 31. | Eubacterium_callanderi.fna |
| 32. | [Eubacterium]_eligens_ATCC_27750.fna |
| 33. | Eubacterium_limosum_strain_ATCC_8486.fna |
| 34. | Eubacterium_limosum_strain_SA11.fna |
| 35. | Lachnoclostridium_phocaeense_strain_Marseille-P3177T.fna |
| 36. | Lachnoclostridium_phytofermentans_strain_KNHs2131.fna |
| 37. | Lachnoclostridium_sp._YL32.fna |
| 38. | Paraclostridium_bifermentans_strain_ATCC_638_plasmid_p14Cb.fna |
| 39. | Paraclostridium_bifermentans_strain_SU1074NT_CbotD0132-U21098_1.fna |
| 40. | Peptoclostridium_acidaminophilum_DSM_3953.fna |
| 41. | Romboutsia_ilealis_strain_CRIB.fna |
| 42. | Romboutsia_sp._Frifi_strain_FRIFI.fna |
| 43. | Romboutsia_sp._MT17.fna |
| 44. | Romboutsia_timonensis_strain_Marseille-P326.fna |
| 45. | Ruminococcus_albus_7_=_DSM_20455.fna |
| 46. | Ruminococcus_bicirculans_strain_803.fna |
| 47. | saccharolyticum_isolate_MGYG-HGUT-02483.fna |

## References

Andrews, S. (2017). FastQC: a quality control tool for high throughput sequence data. 2010.

Antignano, F., Mullaly, S.C., Burrows, K., and Zaph, C. (2011). Trichuris muris infection: a model of type 2 immunity and inflammation in the gut. J Vis Exp. 10.3791/2774.

Aramaki, T., Blanc-Mathieu, R., Endo, H., Ohkubo, K., Kanehisa, M., Goto, S., and Ogata, H. (2019). KofamKOALA: KEGG Ortholog assignment based on profile HMM and adaptive score threshold. Bioinformatics 36, 2251–2252. 10.1093/bioinformatics/btz859.

Atarashi, K., Tanoue, T., Oshima, K., Suda, W., Nagano, Y., Nishikawa, H., Fukuda, S., Saito, T., Narushima, S., Hase, K., et al. (2013). Treg induction by a rationally selected mixture of Clostridia strains from the human microbiota. Nature 500, 232–236. 10.1038/nature12331.

Bach, J.F. (2017). The hygiene hypothesis in autoimmunity: the role of pathogens and commensals. Nat Rev Immunol. 10.1038/nri.2017.111.

Becattini, S., Littmann, E.R., Carter, R.A., Kim, S.G., Morjaria, S.M., Ling, L., Gyaltshen, Y., Fontana, E., Taur, Y., Leiner, I.M., and Pamer, E.G. (2017). Commensal microbes provide first line defense against Listeria monocytogenes infection. J Exp Med 214, 1973–1989. 10.1084/jem.20170495.

Bethony, J., Brooker, S., Albonico, M., Geiger, S.M., Loukas, A., Diemert, D., and Hotez, P.J. (2006). Soil-transmitted helminth infections: ascariasis, trichuriasis, and hookworm. Lancet 367, 1521–1532. 10.1016/s0140-6736(06)68653-4.

Blaser, M.J. (2016). Antibiotic use and its consequences for the normal microbiome. Science 352, 544–545. 10.1126/science.aad9358.

Blaser, M.J., and Falkow, S. (2009). What are the consequences of the disappearing human microbiota? Nat Rev Microbiol 7, 887–894. 10.1038/nrmicro2245.

Bogitsh, B.J., Carter, C.E., and Oeltmann, T.N. (2013). Chapter 16 - Intestinal Nematodes. In Human Parasitology (Fourth Edition), B.J. Bogitsh, C.E. Carter, and T.N. Oeltmann, eds. (Academic Press), pp. 291–327. https://doi.org/10.1016/B978-0-12-415915-0.00016-9.

Bolger, A.M., Lohse, M., and Usadel, B. (2014). Trimmomatic: a flexible trimmer for Illumina sequence data. Bioinformatics 30, 2114–2120. 10.1093/bioinformatics/btu170.

Breitwieser, F.P., Baker, D.N., and Salzberg, S.L. (2018). KrakenUniq: confident and fast metagenomics classification using unique k-mer counts. Genome Biol 19, 198. 10.1186/s13059-018-1568-0.

Brugiroux, S., Beutler, M., Pfann, C., Garzetti, D., Ruscheweyh, H.J., Ring, D., Diehl, M., Herp, S., Lotscher, Y., Hussain, S., et al. (2016). Genome-guided design of a defined mouse microbiota that confers colonization resistance against Salmonella enterica serovar Typhimurium. Nat Microbiol 2, 16215. 10.1038/nmicrobiol.2016.215.

Caballero, S., Kim, S., Carter, R.A., Leiner, I.M., Susac, B., Miller, L., Kim, G.J., Ling, L., and Pamer, E.G. (2017). Cooperating Commensals Restore Colonization Resistance to Vancomycin-Resistant Enterococcus faecium. Cell Host Microbe 21, 592–602 e594. 10.1016/j.chom.2017.04.002.

Carmody, R.N., Sarkar, A., and Reese, A.T. (2021). Gut microbiota through an evolutionary lens. Science 372, 462–463. 10.1126/science.abf0590.

Chaumeil, P.-A., Mussig, A.J., Hugenholtz, P., and Parks, D.H. (2019). GTDB-Tk: a toolkit to classify genomes with the Genome Taxonomy Database. Bioinformatics 36, 1925–1927. 10.1093/bioinformatics/btz848.

Cooper, P., Walker, A.W., Reyes, J., Chico, M., Salter, S.J., Vaca, M., and Parkhill, J. (2013). Patent human infections with the whipworm, Trichuris trichiura, are not associated with alterations in the faecal microbiota. PLoS One 8, e76573. 10.1371/journal.pone.0076573.

Dürre, P. (2014). Physiology and Sporulation in Clostridium. Microbiol Spectr 2, Tbs-0010-2012. 10.1128/microbiolspec.TBS-0010-2012.

Easton, A.V., Quiñones, M., Vujkovic-Cvijin, I., Oliveira, R.G., Kepha, S., Odiere, M.R., Anderson, R.M., Belkaid, Y., and Nutman, T.B. (2019). The Impact of Anthelmintic Treatment on Human Gut Microbiota Based on Cross-Sectional and Pre- and Postdeworming Comparisons in Western Kenya. mBio 10. 10.1128/mBio.00519-19.

Elliott, D.E., and Weinstock, J.V. (2012). Helminth-host immunological interactions: prevention and control of immune-mediated diseases. Ann N Y Acad Sci 1247, 83–96. 10.1111/j.1749-6632.2011.06292.x.

Emms, D.M., and Kelly, S. (2019). OrthoFinder: phylogenetic orthology inference for comparative genomics. Genome Biology 20, 238. 10.1186/s13059-019-1832-y.

Gerritsen, J., Umanets, A., Staneva, I., Hornung, B., Ritari, J., Paulin, L., Rijkers, G.T., de Vos, W.M., and Smidt, H. (2018). Romboutsia hominis sp. nov., the first human gut-derived representative of the genus Romboutsia, isolated from ileostoma effluent. Int J Syst Evol Microbiol 68, 3479–3486. 10.1099/ijsem.0.003012.

Hayes, K.S., Bancroft, A.J., Goldrick, M., Portsmouth, C., Roberts, I.S., and Grencis, R.K. (2010). Exploitation of the intestinal microflora by the parasitic nematode Trichuris muris. Science 328, 1391–1394. 10.1126/science.1187703.

Human Microbiome Project Consortium (2012a). A framework for human microbiome research. Nature 486, 215–221. 10.1038/nature11209.

Human Microbiome Project Consortium (2012b). Structure, function and diversity of the healthy human microbiome. Nature 486, 207–214. 10.1038/nature11234.

Jain, C., Rodriguez-R, L.M., Phillippy, A.M., Konstantinidis, K.T., and Aluru, S. (2018). High throughput ANI analysis of 90K prokaryotic genomes reveals clear species boundaries. Nature Communications 9, 5114. 10.1038/s41467-018-07641-9.

Jenkins, T.P., Rathnayaka, Y., Perera, P.K., Peachey, L.E., Nolan, M.J., Krause, L., Rajakaruna, R.S., and Cantacessi, C. (2017). Infections by human gastrointestinal helminths are associated with changes in faecal microbiota diversity and composition. PLoS One 12, e0184719. 10.1371/journal.pone.0184719.

Kanehisa, M., and Goto, S. (2000). KEGG: kyoto encyclopedia of genes and genomes. Nucleic acids research 28, 27–30. 10.1093/nar/28.1.27.

Kay, G.L., Millard, A., Sergeant, M.J., Midzi, N., Gwisai, R., Mduluza, T., Ivens, A., Nausch, N., Mutapi, F., and Pallen, M. (2015). Differences in the Faecal Microbiome in Schistosoma haematobium Infected Children vs. Uninfected Children. PLOS Neglected Tropical Diseases 9, e0003861. 10.1371/journal.pntd.0003861.

Kernbauer, E., Ding, Y., and Cadwell, K. (2014). An enteric virus can replace the beneficial function of commensal bacteria. Nature 516, 94–98. 10.1038/nature13960.

Kim, Y.G., Sakamoto, K., Seo, S.U., Pickard, J.M., Gillilland, M.G., 3rd, Pudlo, N.A., Hoostal, M., Li, X., Wang, T.D., Feehley, T., et al. (2017). Neonatal acquisition of Clostridia species protects against colonization by bacterial pathogens. Science 356, 315–319. 10.1126/science.aag2029.

Klementowicz, J.E., Travis, M.A., and Grencis, R.K. (2012). Trichuris muris: a model of gastrointestinal parasite infection. Semin Immunopathol 34, 815–828. 10.1007/s00281-012-0348-2.

Kolmogorov, M., Yuan, J., Lin, Y., and Pevzner, P.A. (2019). Assembly of long, error-prone reads using repeat graphs. Nat Biotechnol 37, 540–546. 10.1038/s41587-019-0072-8.

Koyama, K. (2016). Bacteria-induced hatching of Trichuris muris eggs occurs without direct contact between eggs and bacteria. Parasitol Res 115, 437–440. 10.1007/s00436-015-4795-2.

Kupritz, J., Angelova, A., Nutman, T.B., and Gazzinelli-Guimaraes, P.H. (2021). Helminth-Induced Human Gastrointestinal Dysbiosis: a Systematic Review and Meta-Analysis Reveals Insights into Altered Taxon Diversity and Microbial Gradient Collapse. mBio 12, e0289021. 10.1128/mBio.02890-21.

Langmead, B., and Salzberg, S.L. (2012). Fast gapped-read alignment with Bowtie 2. Nat Methods 9, 357–359. 10.1038/nmeth.1923.

Lawson, M.A.E., Roberts, I.S., and Grencis, R.K. (2021). The interplay between Trichuris and the microbiota. Parasitology, 1–8. 10.1017/s0031182021000834.

Lee, S.C., Tang, M.S., Lim, Y.A., Choy, S.H., Kurtz, Z.D., Cox, L.M., Gundra, U.M., Cho, I., Bonneau, R., Blaser, M.J., et al. (2014). Helminth colonization is associated with increased diversity of the gut microbiota. PLoS Negl Trop Dis 8, e2880. 10.1371/journal.pntd.0002880.

Letunic, I., and Bork, P. (2021). Interactive Tree Of Life (iTOL) v5: an online tool for phylogenetic tree display and annotation. Nucleic acids research 49, W293–W296. 10.1093/nar/gkab301.

Loke, P., and Lim, Y.A. (2015). Helminths and the microbiota: parts of the hygiene hypothesis. Parasite Immunol 37, 314–323. 10.1111/pim.12193.

Maizels, R.M. (2020). Regulation of immunity and allergy by helminth parasites. Allergy 75, 524–534. 10.1111/all.13944.

Maizels, R.M., Smits, H.H., and McSorley, H.J. (2018). Modulation of Host Immunity by Helminths: The Expanding Repertoire of Parasite Effector Molecules. Immunity 49, 801–818. 10.1016/j.immuni.2018.10.016.

Maria, R., Dutta, S.D., Thete, S.G., and AlAttas, M.H. (2021). Evaluation of Antibacterial Properties of Organic Gutta-percha Solvents and Synthetic Solvents Against Enterococcus faecalis. J Int Soc Prev Community Dent 11, 179–183. 10.4103/jispcd.JISPCD_422_20.

Martin, I., Djuardi, Y., Sartono, E., Rosa, B.A., Supali, T., Mitreva, M., Houwing-Duistermaat, J.J., and Yazdanbakhsh, M. (2018). Dynamic changes in human-gut microbiome in relation to a placebo-controlled anthelminthic trial in Indonesia. PLoS Negl Trop Dis 12, e0006620. 10.1371/journal.pntd.0006620.

McInnes, L., Healy, J., and Melville, J. (2018). Umap: Uniform manifold approximation and projection for dimension reduction. arXiv preprint arXiv:1802.03426.

Nagano, Y., Itoh, K., and Honda, K. (2012). The induction of Treg cells by gut-indigenous Clostridium. Curr Opin Immunol 24, 392–397. 10.1016/j.coi.2012.05.007.

Price, M.N., Dehal, P.S., and Arkin, A.P. (2009). FastTree: Computing Large Minimum Evolution Trees with Profiles instead of a Distance Matrix. Molecular Biology and Evolution 26, 1641–1650. 10.1093/molbev/msp077.

R Core Team (2021). R: A language and environment for statistical computing. R Foundation for Statistical Computing, Vienna, Austria. https://www.R-project.org/.

Ramanan, D., Bowcutt, R., Lee, S.C., Tang, M.S., Kurtz, Z.D., Ding, Y., Honda, K., Gause, W.C., Blaser, M.J., Bonneau, R.A., et al. (2016). Helminth infection promotes colonization resistance via type 2 immunity. Science 352, 608–612. 10.1126/science.aaf3229.

Ramanan, D., Tang, M.S., Bowcutt, R., Loke, P., and Cadwell, K. (2014). Bacterial sensor Nod2 prevents inflammation of the small intestine by restricting the expansion of the commensal Bacteroides vulgatus. Immunity 41, 311–324. 10.1016/j.immuni.2014.06.015.

Rapin, A., Chuat, A., Lebon, L., Zaiss, M.M., Marsland, B.J., and Harris, N.L. (2020). Infection with a small intestinal helminth, Heligmosomoides polygyrus bakeri, consistently alters microbial communities throughout the murine small and large intestine. Int J Parasitol 50, 35–46. 10.1016/j.ijpara.2019.09.005.

Reynolds, L.A., Smith, K.A., Filbey, K.J., Harcus, Y., Hewitson, J.P., Redpath, S.A., Valdez, Y., Yebra, M.J., Finlay, B.B., and Maizels, R.M. (2014). Commensal-pathogen interactions in the intestinal tract: lactobacilli promote infection with, and are promoted by, helminth parasites. Gut Microbes 5, 522–532. 10.4161/gmic.32155.

Rosa, B.A., Supali, T., Gankpala, L., Djuardi, Y., Sartono, E., Zhou, Y., Fischer, K., Martin, J., Tyagi, R., Bolay, F.K., et al. (2018). Differential human gut microbiome assemblages during soil-transmitted helminth infections in Indonesia and Liberia. Microbiome 6, 33. 10.1186/s40168-018-0416-5.

Sakamoto, M., Iino, T., Yuki, M., and Ohkuma, M. (2018). Lawsonibacter asaccharolyticus gen. nov., sp. nov., a butyrate-producing bacterium isolated from human faeces. Int J Syst Evol Microbiol 68, 2074–2081. 10.1099/ijsem.0.002800.

Sokol, H., Pigneur, B., Watterlot, L., Lakhdari, O., Bermúdez-Humarán, L.G., Gratadoux, J.J., Blugeon, S., Bridonneau, C., Furet, J.P., Corthier, G., et al. (2008). Faecalibacterium prausnitzii is an anti-inflammatory commensal bacterium identified by gut microbiota analysis of Crohn disease patients. Proc Natl Acad Sci U S A 105, 16731–16736. 10.1073/pnas.0804812105.

Song, S.J., Lauber, C., Costello, E.K., Lozupone, C.A., Humphrey, G., Berg-Lyons, D., Caporaso, J.G., Knights, D., Clemente, J.C., Nakielny, S., et al. (2013). Cohabiting family members share microbiota with one another and with their dogs. eLife 2, e00458–e00458. 10.7554/eLife.00458.

Sorbara, M.T., Littmann, E.R., Fontana, E., Moody, T.U., Kohout, C.E., Gjonbalaj, M., Eaton, V., Seok, R., Leiner, I.M., and Pamer, E.G. (2020). Functional and Genomic Variation between Human-Derived Isolates of Lachnospiraceae Reveals Inter- and Intra-Species Diversity. Cell Host Microbe 28, 134–146.e134. 10.1016/j.chom.2020.05.005.

Srivastava, D., Seo, J., Rimal, B., Kim, S.J., Zhen, S., and Darwin, A.J. (2018). A Proteolytic Complex Targets Multiple Cell Wall Hydrolases in Pseudomonas aeruginosa. mBio 9. 10.1128/mBio.00972-18.

Stefka, A.T., Feehley, T., Tripathi, P., Qiu, J., McCoy, K., Mazmanian, S.K., Tjota, M.Y., Seo, G.Y., Cao, S., Theriault, B.R., et al. (2014). Commensal bacteria protect against food allergen sensitization. Proc Natl Acad Sci U S A 111, 13145–13150. 10.1073/pnas.1412008111.

Suchomel, M., Lenhardt, A., Kampf, G., and Grisold, A. (2019). Enterococcus hirae, Enterococcus faecium and Enterococcus faecalis show different sensitivities to typical biocidal agents used for disinfection. J Hosp Infect 103, 435–440. 10.1016/j.jhin.2019.08.014.

Tanizawa, Y., Fujisawa, T., and Nakamura, Y. (2017). DFAST: a flexible prokaryotic genome annotation pipeline for faster genome publication. Bioinformatics 34, 1037–1039. 10.1093/bioinformatics/btx713.

Tee, M.Z., Er, Y.X., Easton, A.V., Yap, N.J., Lee, I.L., Devlin, J., Chen, Z., Ng, K.S., Subramanian, P., Angelova, A., et al. (2022). Gut Microbiome of Helminth Infected Indigenous Malaysians Is Context Dependent. bioRxiv, 2022.2001.2021.477162. 10.1101/2022.01.21.477162.

Tettelin, H., Riley, D., Cattuto, C., and Medini, D. (2008). Comparative genomics: the bacterial pan-genome. Current Opinion in Microbiology 11, 472–477. https://doi.org/10.1016/j.mib.2008.09.006.

Venzon, M., Das, R., Luciano, D.J., Park, H.S., Kool, E.T., Belasco, J.G., Hubbard, E.J.A., and Cadwell, K. (2021). Microbial byproducts determine reproductive fitness of free-living and parasitic nematodes. bioRxiv, 2021.2008.2002.454806. 10.1101/2021.08.02.454806.

Vernikos, G., Medini, D., Riley, D.R., and Tettelin, H. (2015). Ten years of pan-genome analyses. Current Opinion in Microbiology 23, 148–154. https://doi.org/10.1016/j.mib.2014.11.016.

White, E.C., Houlden, A., Bancroft, A.J., Hayes, K.S., Goldrick, M., Grencis, R.K., and Roberts, I.S. (2018). Manipulation of host and parasite microbiotas: Survival strategies during chronic nematode infection. Sci Adv 4, eaap7399. 10.1126/sciadv.aap7399.

WHO (2020). Soil-Transmitted Helminth Infections. WHO Fact Sheets. World Health Organization.

Wickham, H. (2016). ggplot2: Elegant Graphics for Data Analysis (Springer-Verlag New York).

Yoon, S.H., Ha, S.M., Kwon, S., Lim, J., Kim, Y., Seo, H., and Chun, J. (2017). Introducing EzBioCloud: a taxonomically united database of 16S rRNA gene sequences and whole-genome assemblies. Int J Syst Evol Microbiol 67, 1613–1617. 10.1099/ijsem.0.001755.

Zaiss, M.M., Rapin, A., Lebon, L., Dubey, L.K., Mosconi, I., Sarter, K., Piersigilli, A., Menin, L., Walker, A.W., Rougemont, J., et al. (2015). The Intestinal Microbiota Contributes to the Ability of Helminths to Modulate Allergic Inflammation. Immunity 43, 998–1010. 10.1016/j.immuni.2015.09.012.

